# Transposable Element-Gene Splicing Modulates the Transcriptional Landscape of Human Pluripotent Stem Cells

**DOI:** 10.1101/2020.07.26.220608

**Authors:** Isaac A. Babarinde, Gang Ma, Yuhao Li, Boping Deng, Zhiwei Luo, Hao Liu, Mazid Md. Abdul, Carl Ward, Minchun Chen, Xiuling Fu, Martha Duttlinger, Jiangping He, Li Sun, Wenjuan Li, Qiang Zhuang, Jon Frampton, Jean-Baptiste Cazier, Jiekai Chen, Ralf Jauch, Miguel A. Esteban, Andrew P. Hutchins

## Abstract

Transposable elements (TEs) occupy nearly 50% of mammalian genomes and are both potential dangers to genome stability and functional genetic elements. TEs can be expressed and exonised as part of a transcript, however, their full contribution to the transcript splicing remains unresolved. Here, guided by long and short read sequencing of RNAs, we show that 26% of coding and 65% of non-coding transcripts of human pluripotent stem cells (hPSCs) contain TEs. Different TE families have unique integration patterns with diverse consequences on RNA expression and function. We identify hPSC-specific splicing of endogenous retroviruses (ERVs) as well as LINE L1 elements into protein coding genes that generate TE-derived peptides. Finally, single cell RNA-seq reveals that proliferating hPSCs are dominated by ERV-containing transcripts, and subpopulations express SINE or LINE-containing transcripts. Overall, we demonstrate that TE splicing modulates the pluripotency transcriptome by enhancing and impairing transcript expression and generating novel transcripts and peptides.

## Introduction

TEs are repetitive sequences of DNA that make up the majority of mammalian DNA (Jurka et al. 2007; Hutchins and Pei 2015). Whilst the majority of TEs are incapable of retrotransposition, there is growing evidence that inactive incorporated TEs are epigenetically regulated and can alter splicing patterns and generate gene-TE chimeric transcripts that can drive disease and oncogene activity (Jang et al. 2019; Clayton et al. 2020). However, paradoxically, TE expression is also associated with developmental competency and evolutionary innovation (Kunarso et al. 2010; Macfarlan et al. 2012; Fort et al. 2014; Wang et al. 2014; Theunissen et al. 2016; Bourque et al. 2018). TEs are mainly thought to be suppressed by DNA methylation and histone H3K9me3 in somatic cells (Feng et al. 2010; Jonsson et al. 2019), but in the mammalian embryo DNA is demethylated, and human pluripotent stem cells (hPSCs) have reduced epigenetic repression, looser chromatin and a permissive environment for transcript and TE expression (Bulut-Karslioglu et al. 2018). Consequently, TEs are more active during embryogenesis and in hPSCs, compared to somatic cells (Fort et al. 2014), and are expressed in a stage-specifically manner during embryogenesis (Goke et al. 2015; Grow et al. 2015).

In addition to their epigenetic activity, TEs are active genetically and can alter transcript splicing patterns by becoming exonised into transcripts (Macfarlan et al. 2012; Wang et al. 2014; Goke et al. 2015; Grow et al. 2015). TEs make a major contribution to the sequences of long-non-coding RNAs (lncRNAs) (Lev-Maor et al. 2008; Kelley and Rinn 2012; Kapusta et al. 2013; Naville et al. 2016), and whilst lncRNAs have roles in normal biological processes (Goff and Rinn 2015; Bourque et al. 2018; Lu et al. 2020), and disease (Wapinski and Chang 2011; Carlevaro-Fita et al. 2020), the roles of TEs inside the lncRNAs has received less attention. One model suggests that TEs act as functional domains within lncRNAs, something akin to globular domains of proteins (Johnson and Guigo 2014; Chishima et al. 2018). However, the full contribution of TEs to transcript splicing in both the normal and disease state remains unexplored. Interestingly, the splicing of TEs into normal pluripotency genes has been observed in cancer (Jang et al. 2019), suggesting TE-splicing can contribute to human disease. Consequently, it is important to understand how TEs contribute to the coding and non-coding transcriptome (You et al. 2017; Ma et al. 2019; Morillon and Gautheret 2019), if hPSCs are to be used in cell replacement therapy (Fort et al. 2014; Schumann et al. 2019). However, due to limitations in short-read based sequencing, accurate transcriptome maps have been elusive. Much of this is due to the relatively low expression of the TEs and lncRNAs (Lagarde et al. 2017), the repetitive nature of TEs, and problems in using short-reads to assemble longer transcripts (Steijger et al. 2013; Babarinde et al. 2019).

To explore the splicing patterns of TEs in the normal, non-diseased, state, we took advantage of the large number of hPSC short-read-RNA-seq samples, which we supplemented with long-read-RNA-seq to build a map of the gene-TE transcriptome of hPSCs. Our analysis shows that TEs are expressed and spliced into both coding and non-coding RNAs, and had strong effects on gene expression. TEs make up a substantial fraction of the non-coding transcript complement, and they also alter the peptide output of the cell, producing fragments of endogenous viral proteins and small peptides. Finally, using single-cell RNA-seq we show that the hPSC-state is dominated by HERVH and LTR7-containing transcripts, which are associated with proliferating cells and are rapidly lost on differentiation as SINE and LINE-containing transcripts become dominant.

## Results

### Transcript assembly from short and long reads

To build a hPSC transcriptome, we started with 197 publicly available short read (SR) paired-end-only hPSC RNA-seq data samples. To quality control the data, we mapped the hPSC samples to the hg38 genome assembly using HISAT2 (Kim et al. 2019). Only samples with a minimum mapping rate of 70% were retained, leaving 171 RNA-seq samples (**Supplemental Fig. S1A**). There are widespread errors in metadata annotation of publically available transcriptome data (Toker et al. 2016), consequently, to ensure that the samples were undifferentiated hPSCs, we measured gene-level expression (Hutchins et al. 2017). hPSC samples were removed if they were outliers in a co-correlation analysis, or did not express hPSC-marker genes, or expressed differentiation-specific genes (**Supplemental Fig. S1B and C**). This left 150 samples that passed our quality control metrics (**Supplemental Table S1, and Supplemental Fig. S1D**). We used the 150 confirmed hPSC samples to assemble transcripts using HISAT2/StringTie (Pertea et al. 2015; Kim et al. 2019) (**Supplemental Fig. S1E**). In total ~13 billion reads, with a median mapped fragment size of 225 bp of which 92% could be aligned to the hg38 genome assembly. StringTie (Pertea et al. 2015) assembled an initial set of 279,051 transcripts, which was reduced to 272,268 after removing transcripts shorter than 200 bp (**Supplemental Fig. S1F)**. This dataset comprised the transcripts assembled from the SR data.

Transcript assembly from SR can be problematic (Wang et al. 2019), especially if the transcripts contain transposable elements (Babarinde et al. 2019). This is reflected in the large number of single-exon fragments from the SR assembly (72,902; 27%). Although some of these fragments might be genuine single-exon transcripts, many are likely to be fragmentary assemblies from SR. To address this, we augmented our assembly with SMRT long-read (LR) sequencing using the PacBio platform. We sequenced the hESC cell line H1 and the iPSC cell line S0730 in duplicate (Zhou et al. 2011). Transcripts were identified from long reads using the isoseq pipeline, and were aligned to the genome using GMAP (Wu and Watanabe 2005) (**Supplemental Fig. S1E**). Long read assembly produced 53,168 unique transcripts that were at least 200bp long (**Supplemental Fig. S1F**).

Both SR-based and LR-based transcriptomes have disadvantages: SR-based assemblies tend to be unreliable (Steijger et al. 2013), and LR-based assemblies can detect rare transcripts and are poor at quantifying expression. Consequently, to arrive at a transcriptome representation, we set two expression level conditions that a transcript must pass: (1) Expression >=0.1 TPM (transcripts per million) in at least 50 of the samples. (2) The average per-base coverage reported by StringTie must be greater than 1 for all exons. Additionally, we also deleted single-exon transcripts that appear to be derived from incomplete splicing, if they shared edges near perfectly with an annotated intron (**Supplemental Fig. S1G**). The final assembly contained 101,492 transcripts, of which 13,177 (13%) were single-exons. Using these criteria, 71% of the LR transcripts were retained, but only 17% of the SR transcripts were kept in the final assembly (**Supplemental Fig. S1F, H**). To validate our assembly pipeline, we designed RT-PCR primers spanning an intron (**Supplemental Table S2**) and could detect 31 out of 40 transcripts including 8 out of 9 long-read transcripts (**Supplemental Fig. S2A**). We confirmed the accuracy of the 5’ ends of our transcript assembly using iPSC and hESC deepCAGE data (Consortium et al. 2014) (**Fig. 1A, S2B**). There was a linear relationship between deepCAGE expression and RNA-seq TPM for the transcripts we could unambiguously assign a deepCAGE cluster (**Supplemental Fig. S2C**). Our final transcript assembly contained 58,201 transcripts with SR support, 6,881 with LR support and 36,410 with both SR and LR support (**Fig. 1B and Supplemental Table S3**).

**Figure 1.**
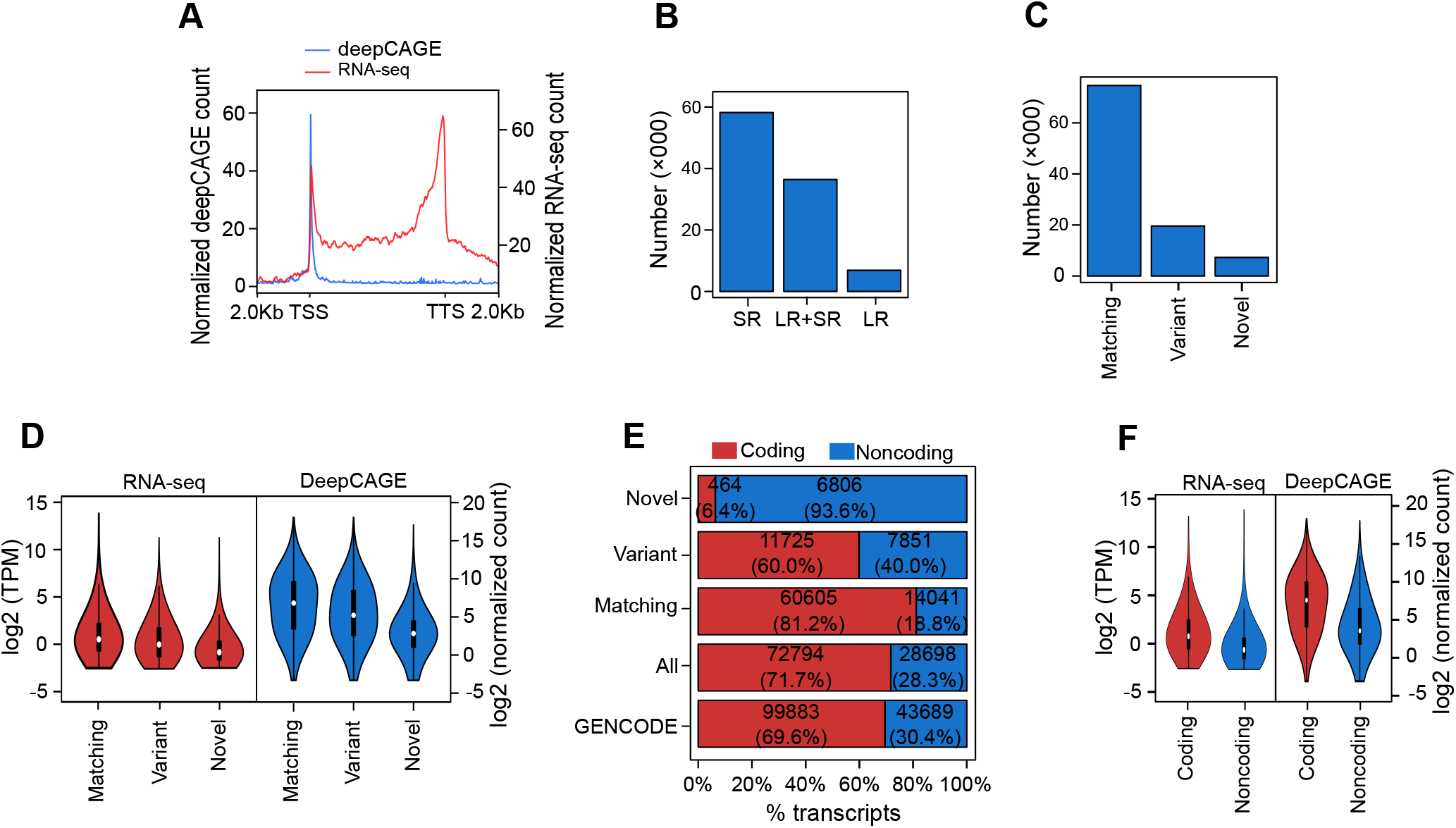
Combining short-read RNA-seq and long read RNA-seq to assemble a high-quality hPSC transcriptome. (A) Pileup of deepCAGE and RNA-seq reads across the lengths of the transcripts from TSS to TTS and the flanking 2 kb regions. (B) Number of transcripts (thousands) supported by short-read (SR), long read (LR) only, or both (SR+LR). (C) Number of transcripts defined as matching (internal exon structure matches exactly to a GENCODE transcript), variant (shares any exon or overlapping exon segment with a GENCODE transcript) or novel (does not share any exon base pair with a GENCODE transcript). (D) Violin plots showing RNA-seq log2 transcripts per million (TPM) and deepCAGE log2 normalized tag count expression for all unambiguous TSSs in the hPSC assembled transcriptome. (E) Percentage of coding and non-coding transcripts by transcript class. (F) Expression level of coding and non-coding transcripts, for short-read RNA-seq (left violins) or deepCAGE data (right violins).

### Identification of novel transcripts and isoforms

We next classified the hPSC transcripts based on the GENCODE transcriptome. We defined three categories: Matching (all internal splices match a GENCODE transcript and at least 75% total exon length overlap); Variant (with an exon overlapping any exon from a GENCODE transcript but not completely matching), and Novel (with no exon overlapping any GENCODE exon) (**Fig. 1C**). Whilst most transcripts shared a splicing pattern with a GENCODE transcript, 26,846 transcripts were variant or novel (**Fig. 1C**). For each assembled transcript, we defined completeness as the percent of exons or splices of the closest GENCODE transcript that were correctly retrieved (**Supplemental Fig. S2D**). Transcripts supported by both LR and SR tend to have higher GENCODE exon and splice completeness (**Supplemental Fig. S2E**), although there remains a sizeable number of transcripts supported only by SR, suggesting our LR data set has not fully saturated the transcriptome (**Supplemental Fig. S2F**). The known transcripts tended to be expressed more strongly than novel transcripts, as measured by both RNA-seq and deepCAGE (**Fig. 1D**).

### A census of coding and non-coding transcripts enriched in pluripotent stem cells

FEELnc was used to discriminate coding and non-coding transcripts (Wucher et al. 2017), of which 28,699 (28%) were non-coding, and the majority of novel transcripts were non-coding (**Fig. 1E, and Supplemental Fig. S2G-H**). Conversely, the majority (~81%) of the GENCODE-matching transcripts were protein-coding. As expected, the expression of lncRNAs was lower than protein coding transcripts (**Fig. 1F, and Supplemental Fig. S2I**). Genes can be expressed in both a cell type-independent and cell type-specific manner (Hutchins et al. 2017). To identify the transcripts important for hPSCs function, we measured the expression of our transcript assembly using the BodyMap dataset (Derrien et al. 2012), and divided the transcripts into hPSC-enriched, -nonspecific or -depleted, based on the Z-score against the BodyMap expression dataset (**Fig. 2A, and Supplemental Fig. S2J**). This approach could recover known hPSC-enriched transcripts such as *NANOG*, *POU5F1* and *SALL4* (**Fig. 2B**). There was no specific bias in coding and non-coding transcripts between the hPSC expression categories (**Fig. 2C**). However, novel transcripts were more likely to be enriched in hPSCs (**Supplemental Fig. S2K**). Finally, non-coding transcripts had a lower level of expression compared to coding transcripts across all categories (**Fig. 2D**). Using LR and SR we have generated an accurate transcriptome for hPSCs, that describes known and novel transcripts, coding and non-coding, and hPSC-enriched and -depleted transcripts. We will use this transcriptome to explore the effect of TEs.

**Figure 2.**
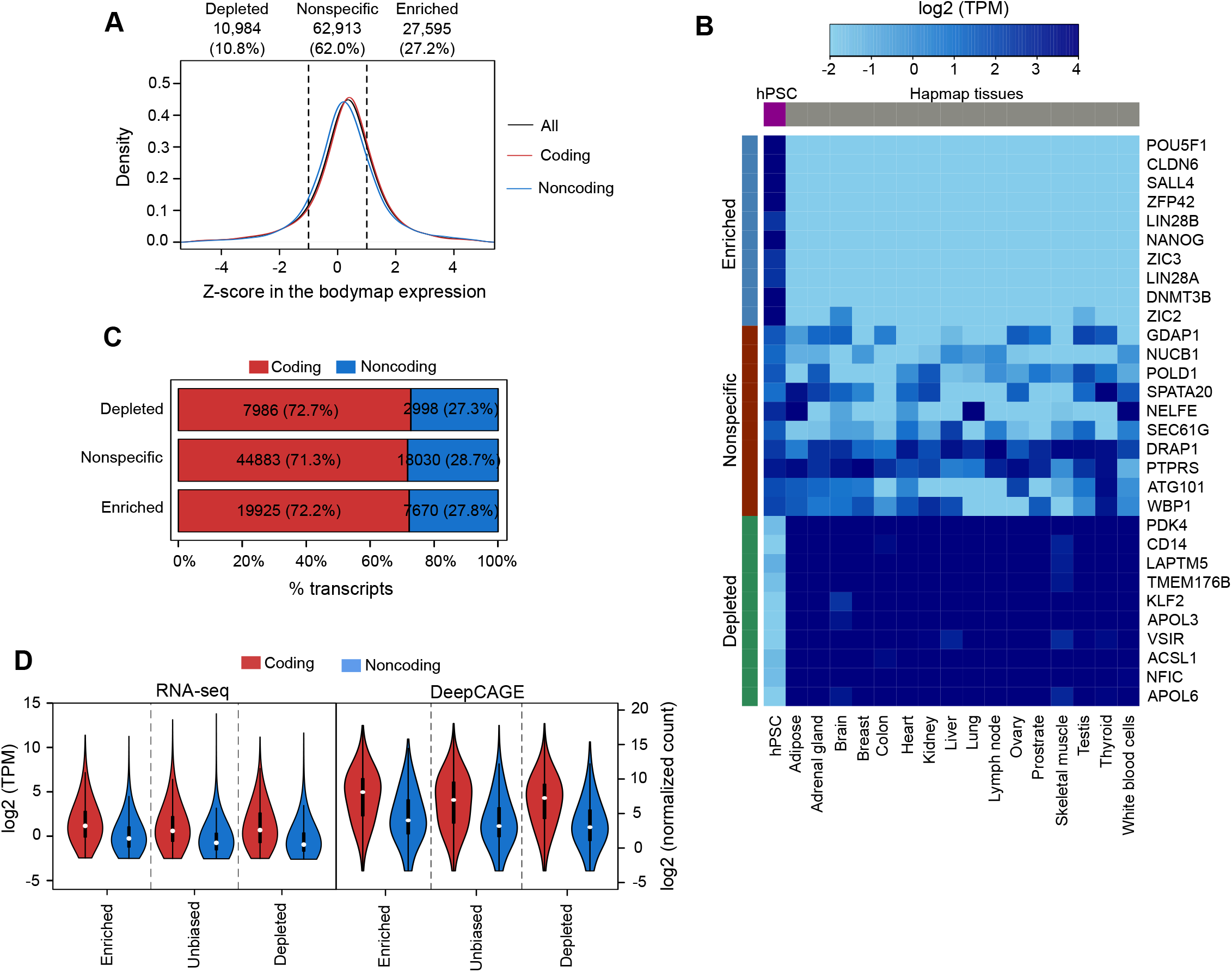
Transcript dynamics of the hPSC transcriptome. (A) Z-score of all, coding, and non-coding transcripts against the Human BodyMap 2.0 dataset (Derrien et al. 2012). (B) Heatmap showing expression of selected hPSC-enriched, -nonspecific and -depleted transcripts in hPSCs and the BodyMap samples. (C) Percent of the coding and non-coding transcripts in the indicated hPSC expression categories. (D) Violin plots showing the RNA-seq expression levels (left panel; TPM: transcripts per million) or deepCAGE data (right panel; in normalized read counts) for the indicated expression classes for coding and non-coding transcripts.

### TEs are widely spliced into protein coding genes and modulate expression levels

TEs are found in the untranslated regions (UTRs) of coding genes (Bourque et al. 2018), and contribute to lncRNAs (Kelley and Rinn 2012; Kapusta et al. 2013). Additionally, deepCAGE data has revealed pluripotent-specific TSSs (transcription start sites) originating inside TEs (Faulkner et al. 2009; Fort et al. 2014). We searched our assembled transcriptome using nhmmer (Wheeler and Eddy 2013), using the DFAM models of TEs (Hubley et al. 2016), and identified 37,493 transcripts (37%) that contained at least one TE (**Supplemental Table S4**). About 22% of the matching transcripts contained a TE, which was similar to GENCODE, however the variant coding transcripts had a higher percentage of TE-containing transcripts in hPSCs (**Fig. 3A**). There was a very small number of novel coding transcripts (411), of which 139 (34%) were predicted to contain a TE. As this number is close to the false positive rate to detect coding transcripts, we do not explore these further (**Fig. 1E, Supplemental Fig. S2G, and Supplemental Table S3**). Surprisingly, hPSC-enriched transcripts were less likely than hPSC-nonspecific or hPSC-depleted transcripts to contain a TE (**Fig. 3B**). This effect was not unique to the subset of novel transcripts assembled by us, as the same pattern is observed for transcripts matching the GENCODE annotations (**Fig. 3C**). This is unexpected as TEs are thought to be more actively expressed in the epigenetically relaxed hPSCs.

**Figure 3.**
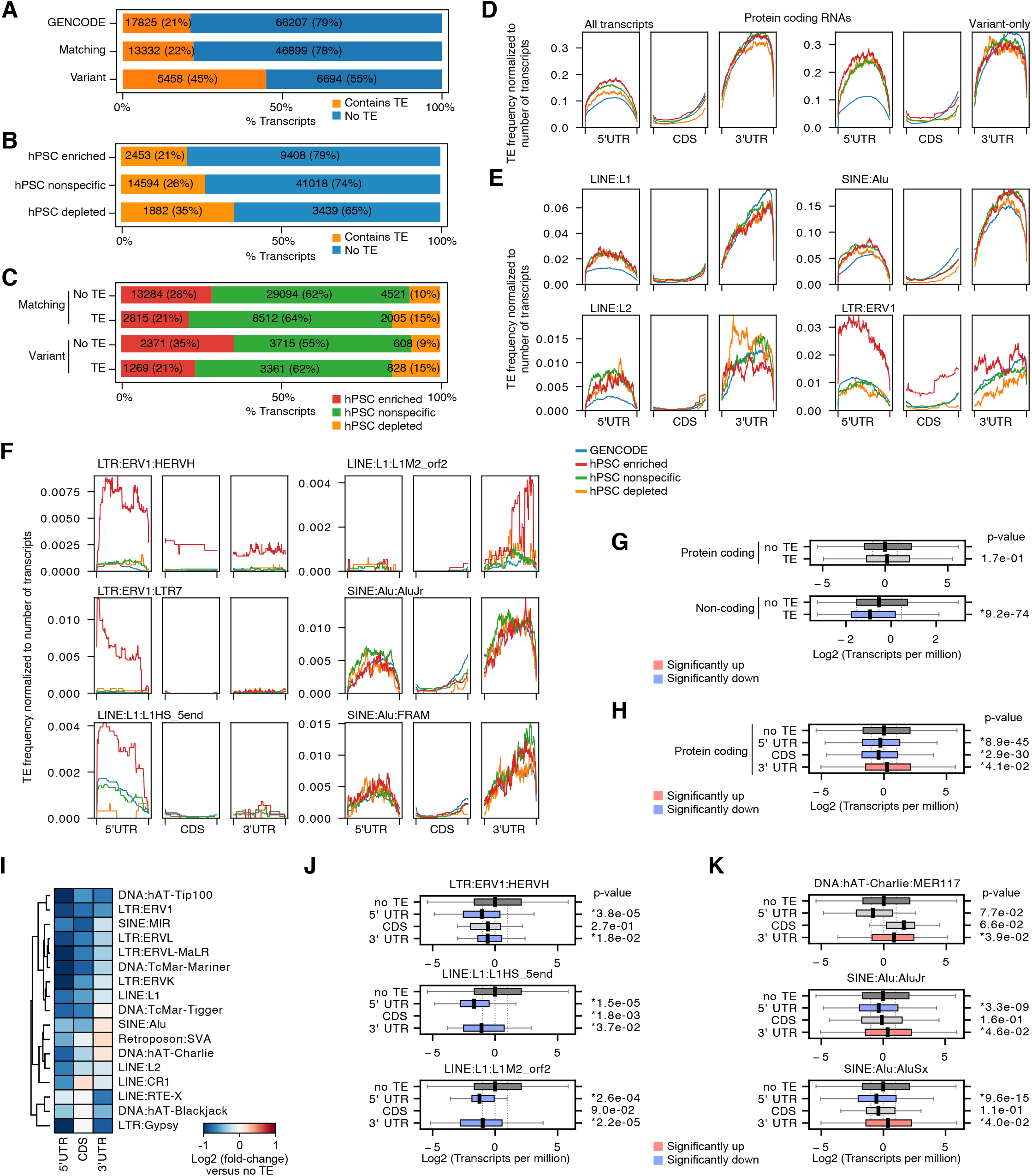
TEs spliced into coding sequence genes remodel the expression dynamics of pluripotent stem cells. (A) Proportion of protein coding transcripts in the GENCODE database, of hPSC-expressing transcripts matching (to GENCODE), or variant (share transcript structure with a known transcript), and the proportions that contain at least 1 TE or no TEs. (B) Proportion of coding transcripts that are enriched, nonspecific or depleted in hPSCs (C) Proportions of matching or variant transcripts with or without a TE that are hPSC-enriched, - nonspecific or -depleted. (D) Relative density of all TE insertions in all or variant transcripts’ 5’ UTR, CDS or 3’UTR. Each plot is normalized to the length of the transcript region. The blue line indicates all GENCODE transcripts, the red, green and yellow lines indicate hPSC-specific expression categories. (E) Density plots (as in panel D) for LINE L1, L2, SINE Alu, and LTR ERV1 TEs. (F) Density plots for the indicated LTR, LINE and SINE types. (G) Box plot for the expression levels of protein coding and non-coding transcripts with or without a TE in its sequence. p-value is from a Welch’s t-test versus no TE-containing transcripts. (H) Box plot for the expression levels of protein coding transcripts with or without a TE in the 5’UTR, CDS or 3’UTR. p-value is from a Welch’s t-test versus no TE-containing transcripts. (I) Heatmap showing the fold-change of transcripts containing one or more of the indicated subtypes of TE versus no TE-containing transcripts. (J) Box plot for the expression levels of the transcripts containing the indicated TEs in the 5’UTR, CDS or 3’UTR. p-value is from a Welch’s t-test versus no TE-containing transcripts. (K) As in panel J, but for example transcripts with TEs in the 3’UTR that lead to higher levels of expression. p-value is from a Welch’s t-test versus no TE-containing transcripts.

We next analyzed TE insertion frequency into the UTRs or CDS of coding transcripts (**Fig. 3D**). We annotated the CDS from either the GENCODE reference or using the longest open reading frame (ORF). There was an enrichment of TEs in the UTRs, particularly inside the 3’UTRs, and a depletion in the CDS (**Fig. 3D**). We next asked whether integration sites vary across TE families. LINE:L1, L2 and SINE:Alu elements were specifically enriched in both UTRs, but particularly in the 3’UTRs (**Fig. 3E, and Supplemental Fig. S3A, S3B**). LINEs and SINEs were robustly depleted in the CDS, compared to the UTRs (**Fig. 3E**), and most LINE:L1 TEs were biased to the UTRs (**Fig. 3E**). For example, LINE:L1:L1M2_orf2 was strongly biased to the 3’UTR, and LINE:L1:L1HS_5end was specifically enriched in the 5’UTR (**Fig. 3F, and Supplemental Fig. S3C**). Intriguingly, the L1HS family of LINEs are active and capable of retrotransposition in humans (Beck et al. 2010), and their presence here in the 5’UTRs of genes suggests activity in hPSCs.

In a pattern similar to LINEs, several SINE family members were spliced into the 3’ ends of transcripts (**Fig. 3E, and Supplemental Fig. S3B**), whilst HERVH/LTR7 TEs were specifically enriched in the 5’UTR (**Fig. 3F**). Previous analysis of deepCAGE data showed that HERVH/LTR7 acts as a hPSC-specific transcription start site (Fort et al. 2014). Our assembly included previously reported LTR7/HERVH initiating transcripts (Wang et al. 2014) (**Supplemental Fig. S3D**). Additionally, we could assemble transcripts with HERVH spliced throughout the transcripts, including the CDS and 3’UTR (**Fig. 3F, and Supplemental Fig. S3E**), indicating that HERVH can function as an exonised unit. LTR7, the HERVH-accompanying LTR, was restricted to only the 5’UTR (**Fig. 3F**).

We next looked at the effect of TE insertions on transcript expression. Non-coding transcripts containing TEs had significantly lower expression levels, however coding transcripts were overall unaffected by the presence of a TE (**Fig. 3G**). It was previously reported that retrotransposons in the 3’UTR lead to decreased isoform expression (Faulkner et al. 2009), hence we explored if TE location within a coding transcript influences expression. Transcripts with a TE in the 5’UTR or CDS had significantly lower expression, however those with a TE inside the 3’UTR were significantly (though modestly) up-regulated (**Fig. 3H**). We next looked at the individual subtypes of TEs, and found that most subtypes of TE led to a lower level of expression (**Fig. 3I**). Transcripts with HERVHs, or L1 LINEs had significantly lower expression no matter the location of their insertion (**Fig. 3I, J**). However, some TEs could be inserted into the 3’UTR, and expression would be higher. For example, SINEs and the DNA:hAT-Charlie family (**Fig. 3I, K**). Overall, the effect of TEs in transcripts is both transcript context and TE-family dependent, and whilst LTRs tend to be uniformly deleterious for expression, some SINEs and DNA transposons in the 3’UTR are associated with enhanced expression.

### TE splicing affects the proteome of pluripotent cells

We next analyzed whether TE insertions alter the amino acid sequences of coding transcripts. We divided TE insertions into classes based upon the TE insertions effect on the CDS, and counted the number of TE insertions (**Fig. 4A**). We did not detect any in-frame TE insertions and found only 63 frame-shift insertions (**Fig. 4A**). A sizeable number of TE insertions led to a premature STOP or to the introduction of a new ATG that would disrupt the pre-existing CDS. However, the most common type of TE insertion caused a disruption or frame-shift that led to a longer CDS becoming the most likely predicted CDS (**Fig. 4A**). These transcripts are predicted as coding, however they may not yield a peptide, as they may lack productive translation or peptide products may be rapidly degraded. Additionally, some TE types, particularly the SINE/Alu and SVA TEs, often have coding-like signatures, but do not encode proteins (Jungreis et al. 2018). To estimate how many of these TE modified CDSs are translated, we extracted the peptide sequences of the putative CDS, deleted any regions that had a >95% BLAST hit against any GENCODE protein, removed peptides shorter than 20 amino acids and retained only those that had at least one Trypsin/Lys-C cleavage site. These strict criteria mean that we would only detect peptides derived from the impact of the TE, or from TE-derived sequences. We then used the HipSci proteomics LC-MS/MS data (Kim and Pevzner 2014; Kilpinen et al. 2017) to search for peptides. Overall, we could detect at least one peptide match for ~17-20% of transcripts (**Fig. 4B and Supplemental Table S5**), indicating that at least some TE-modified transcripts are translated (**Supplemental Fig. S4A**). The majority of peptides were not derived directly from the TE, but correspond to novel ORFs outside the TE sequence (**Fig. 4C**). Yet, a small number of peptides are directly encoded by TEs (**Fig. 4D**). Many of the peptides corresponded to *gag*, *pol* and *env* but not *pro*, of a putative progenitor HERVK (Dewannieux et al. 2006) (23 unique peptides in total) (**Supplemental Fig. S4B**). These results are consistent with reports that some human cells synthesize viral-like structures, particularly in the placenta (Kammerer et al. 2011), and HERVK RNAs and proteins are expressed in hPSCs (Fuchs et al. 2013). Our data suggests that these viral proteins are derived from up to nine transcripts (**Supplemental Table S5**). Interestingly, we find evidence for coding potential for a variant hPSC-enriched transcript *PCAT14* (prostate cancer-associated transcript 14) that has hitherto been considered non-coding. We observed 13 unique HERVK peptides derived from PCAT14 that encode a near-complete product of *gag* and fragments of *pol* (**Fig. 4E, Supplemental Fig. S5A-C and Supplemental Table S5**). We did not observe any peptides from HERVH, in agreement with a previous report that the ORFs are not functional (Santoni et al. 2012). In addition to HERVs, we also detected peptides derived from SINEs, which was unexpected as SINEs do not encode proteins and rely on LINEs for retrotransposition. Using BLAST to search for matching peptides against the human non-redundant protein set produced no significant hits. In summary, these data indicate that TE insertions modulate the coding transcriptome of pluripotent cells, and can give rise to novel peptides and the ectopic expression of viral proteins.

**Figure 4.**
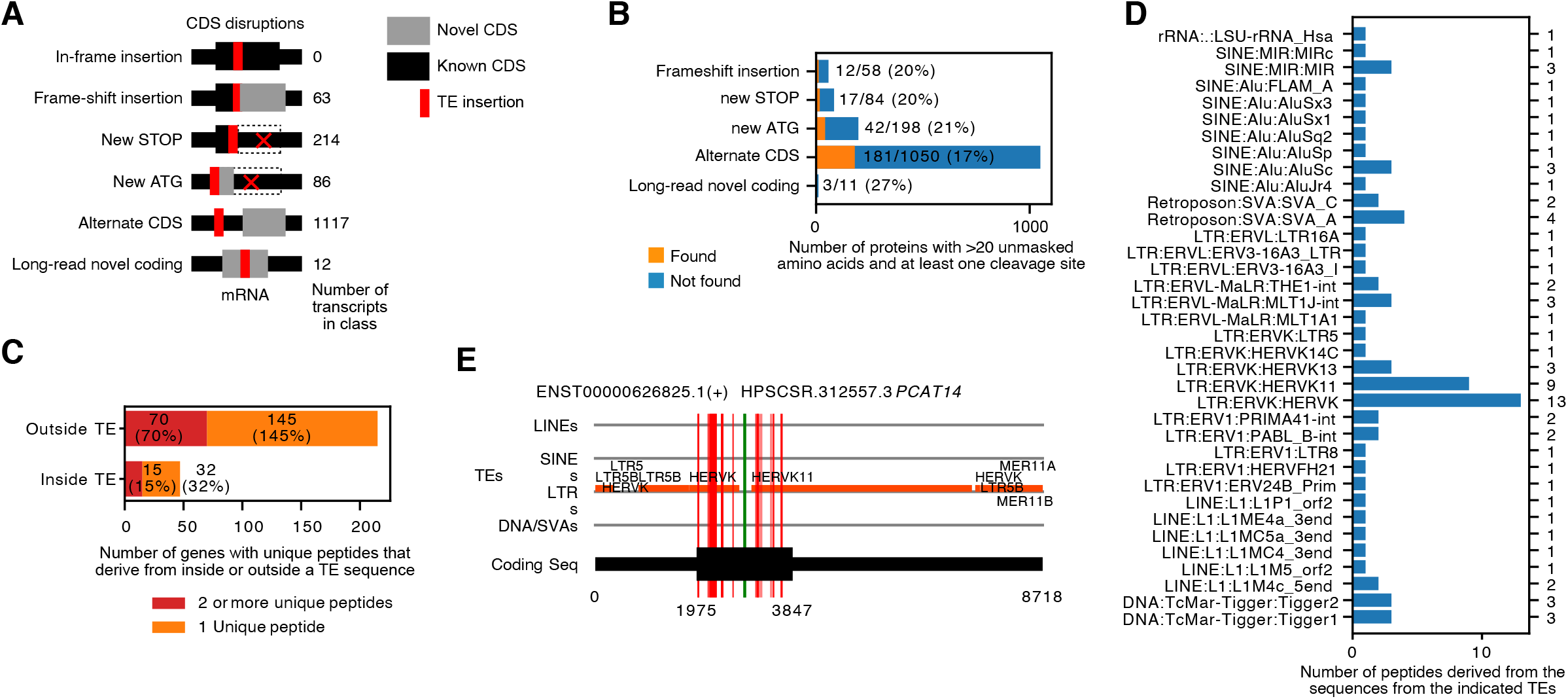
TEs disrupt coding sequences and in limited instances code for peptides. (A) Schematic of the type of TE insertions into the coding sequences, and the number of transcripts in each class. Grey regions indicate novel CDS caused by the TE insertion, black indicates the known CDS, dotted lines indicate where the CDS would be if the TE was not present, and red indicates the TE insertion. Transcripts were assigned to each category from top to bottom by classification. (B) Number of novel CDS peptide fragments that can be detected in the HipSci mass spectrometry dataset. A protein was considered found if at least 1 unique peptide mapped to it. (C) Number and percentage of transcripts with a unique peptide derived from inside or outside the TE sequence. (D) Numbers of unique peptides derived from each type of TE. (E) Example domain map for *PCAT14*, a variant transcript enriched in hPSCs. The location of TEs in the LINE, SINE LTR or DNA/SVAs is indicated on the horizontal lines. The coding sequence is indicated using the thick black bar. The red vertical bars indicate peptides that map inside a TE, the green vertical bar indicates a peptide mapping outside a TE.

### TEs wire the lncRNA complement of hPSCs

Previous reports show that TEs constitute a major part of lncRNAs (Kelley and Rinn 2012; Kapusta et al. 2013). Consistently, we observed a large number of non-coding transcripts containing at least 1 TE (65%), slightly less that the 83% reported in a smaller set of lncRNAs (Kelley and Rinn 2012). Of the transcripts matching GENCODE, 45% had a TE, whilst the variant and novel transcripts were more likely to contain a TE (**Fig. 5A**). Interestingly, and in contrast to coding transcripts, novel and variant non-coding transcripts containing a TE were more likely to be enriched in hPSCs (**Fig. 5B**).

**Figure 5.**
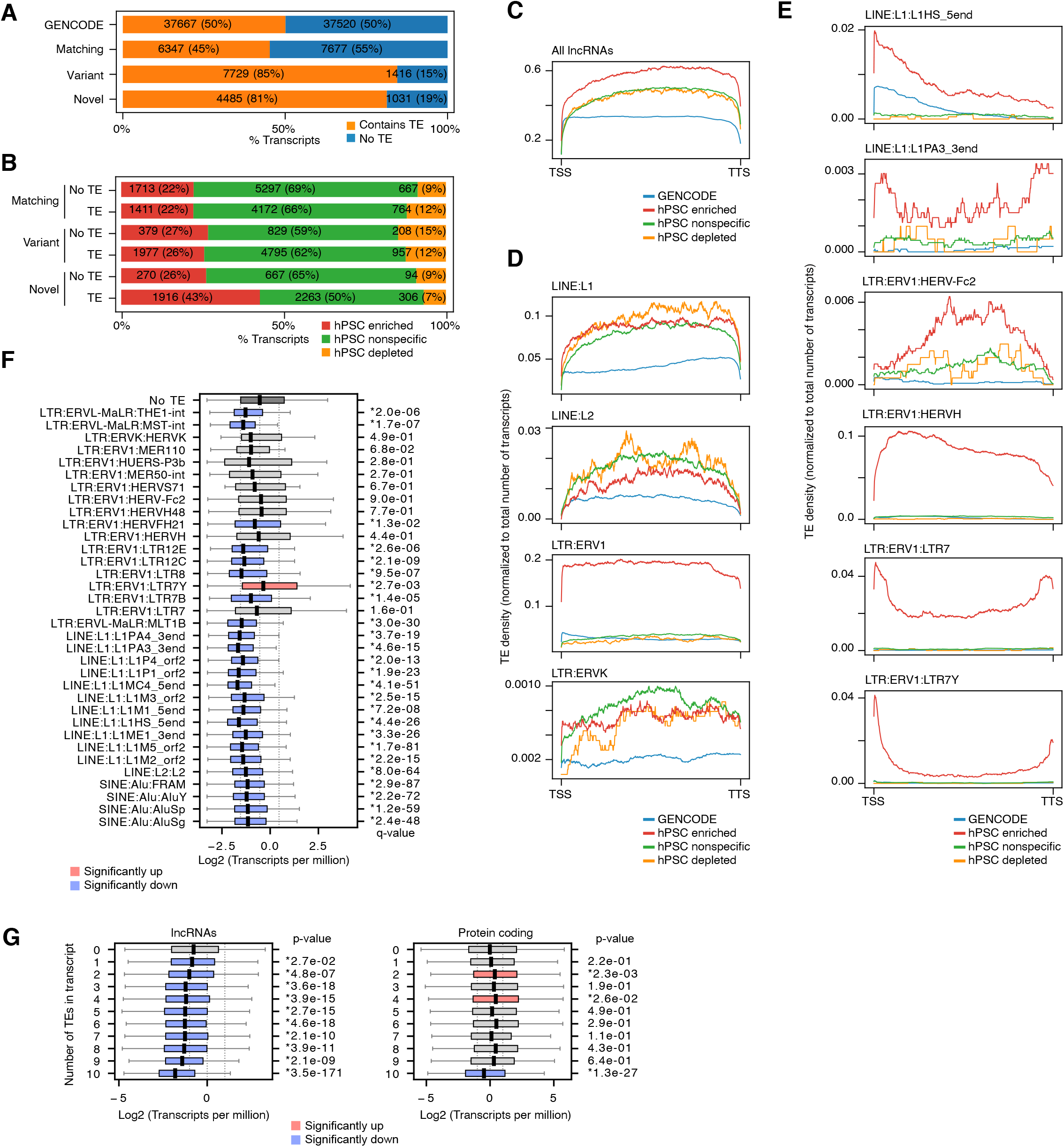
Widespread type-specific splicing of TEs into hPSC-specific lncRNAs. (A) Frequency of TE insertions in GENCODE, and hPSC-matching, variant and novel lncRNAs. (B) Frequency of hPSC expression classes for matching, novel and variant lncRNAs, including or excluding a TE. (C) Frequency of TEs across the normalized length of all lncRNAs. TSS=Transcription start site; TTS=Transcription termination site. Frequency is normalized to the total number of transcripts in each expression class. (D) Examples frequency showing the insertion density frequency for LINE L1, L2 and ERV1 and ERVK TEs. (E) As in panel C and D, except showing selected TE types. (F) Box plots showing the expression of lncRNAs containing the indicated TE types, versus all transcripts with no TE. q-value is from a Welch’s t-test with Bonferroni-Hochberg multiple test correction (for all families of TE). Red boxplots indicate significantly up, blue indicates significantly down, grey no significant change. (G) Box plots showing the effect of multiple TE insertions on non-coding and protein-coding transcripts. p-value is from a Welch’s t-test versus transcripts containing zero TEs.

We next looked at the position of TEs inside lncRNA sequences. TEs are spliced into lncRNAs, and are found anywhere from the TSS to the TTS (transcription termination site) with a slight bias towards the 3’ end that was more pronounced in hPSC-enriched transcripts (**Fig. 5C**). Interestingly, compared to GENCODE, hPSC transcripts were more enriched with TEs, and hPSC-enriched transcripts contained even more TEs (**Fig. 5C, 5D, and Supplemental Fig. S6A**). This indicates that the contribution of TEs to non-coding transcripts is increased in hPSCs. At the TE family level, LINEs were enriched in lncRNAs, such as LINE:L1:L1PA3_3end and LINE:L1:L1Hs_5end (**Fig. 5E, and Supplemental Fig. S6B**). The L1HS family are active and capable of retrotransposition in humans, and although we did not observe intact L1HS sequences or peptides, that L1HS TE fragments are contained inside lncRNAs suggests they are under active epigenetic regulation (**Fig. 3F and 5E**). In addition to LINEs, the SINEs, DNA, SVA and Satellite type repeats/TEs were also enriched (**Supplemental Fig. S6A, S6C-E**). However, overall, the LTRs had the most complex patterns of splicing (**Fig. 5E**), and especially the ERV1 and ERVK families of TEs (**Supplemental Fig. S6F, S6G**). LTR7, the LTR for HERVH, can function as a hPSC-specific TSS (Fort et al. 2014). However, unlike coding transcripts, LTR7 not was biased to the TSS and there was splicing of LTR7 and HERVH along the entirety of the lncRNAs (**Fig. 5E**). There were similarly complex patterns of other LTRs, for example HERVFH21 and HERVH48 were found in the middle of transcripts, and not at the TSS or TTS (**Supplemental Fig. S6G**). Overall, these results indicate that TEs are more likely to be spliced into hPSC-enriched non-coding transcripts, which is in contrast to hPSC-enriched coding transcripts which are depleted of TEs (**Fig. 3A, 3B and 5B**).

Finally, we looked at the effect of TE insertions on lncRNA expression. In contrast to coding transcripts, the presence of a TE inside the lncRNA was more likely to be deleterious for expression (**Fig. 3G**). Many of the hPSC-specific transcripts with TEs had significantly reduced expression compared to transcripts containing no TE (**Fig. 5F**). Additionally, multiple TE insertions resulted in decreased expression, an effect not seen in coding transcripts (**Fig. 5G**). These results indicate that in direct contrast to coding transcripts TE insertions are deleterious for non-coding transcript expression.

### Single cell RNA-seq expression of lincRNAs, heterogeneity in TE splicing

We took advantage of our hPSC-specific high-quality transcript assembly to look at single cell RNA-seq (sc-RNA-seq) data. Sc-RNA-seq technology can measure specific expression of genes in individual cells. However, transcript identity can be ambiguous as the reads are heavily biased to the 3’ ends. Consequently, we reduced the transcripts to those with unique non-overlapping strand-specific 3’ ends, and collapsed overlapping transcript 3’ends to a single transcript. This resulted in a total set of 88,520 transcripts (87% of the total superset of transcripts). We generated two sc-RNA-seq datasets, one from WIBR3 hESCs and another from S0730/c11 iPSCs (Zhou et al. 2011), supplemented with 5 samples from WTC iPSCs from E-MTAB-6687 (Nguyen et al. 2018), and two UCLA1 line hESC samples from GSE140021 (Chen et al. 2019). We aligned the reads to the hg38 genome assembly and annotated the reads to our 3’end transcript database. On average 50-70% of the reads could be aligned to this 3’ end transcriptome (in comparison 60-70% align to GENCODE for the same samples). The majority of the 3’ ends show strand-specific read pileups (**Supplemental Fig. S7A**) indicating that our transcript assembly has accurate 3’ ends and can be used for scRNA-seq. After filtering and normalization, we retrieved 30,001 cells, and 35,400 transcript ends. Projection of the cells into a UMAP (Uniform Maniform and Approximation Projection) plot showed the hESCs and iPSCs mixed well, and there was no bias in the replicate samples (**Supplemental Fig. S7B, S7C**). We could detect five major clusters of cells (**Fig. 6A**). GO (gene ontology) analysis for differentially regulated transcripts specific to each cluster and marker genes expression indicated that all clusters except clusters 3 and 4 were pluripotent (**Fig. 6B, 6C, and Supplemental Fig. S7D**). Analysis of the expression of G1, S and G2/M-specific genes indicated that clusters 0 and 1 had higher expression of G2/M and proliferative genes (**Fig. 6D, E**), suggesting these are highly proliferative cells identified previously (Nguyen et al. 2018; Lau et al. 2020), positive for *EPCAM*, *CD9* and *PODXL* (GCTM2) (**Fig. 6E, and Supplemental Fig. S7E, S7F**). Conversely, clusters 2 and 4 showed reduced levels of G2/M (**Fig. 6D**). Intriguingly, our data contained two novel clusters of cells (cluster 3 and 4), that was predominantly made up of hESCs, rather than iPSCs (**Supplemental Fig. S7B**). GO analysis suggested that these cells were undergoing differentiation (**Fig. 6B**). Marker genes for a specific lineage were not evident, but there was a strong shift in epithelial and mesenchymal genes, as *CDH1* and *EPCAM* were downregulated and *CDH2* and *VIM* were up-regulated (**Supplemental Fig. S7E, S7G**). This is reminiscent of the epithelial-mesenchymal transition we saw previously in the early stage of hepatocyte differentiation (Li et al. 2017). The main difference between the two clusters was in cell cycle activity, as many cells were in G2/M in cluster 3, but cluster 4 showed reduced cell cycle activity (**Fig. 6D**).

**Figure 6.**
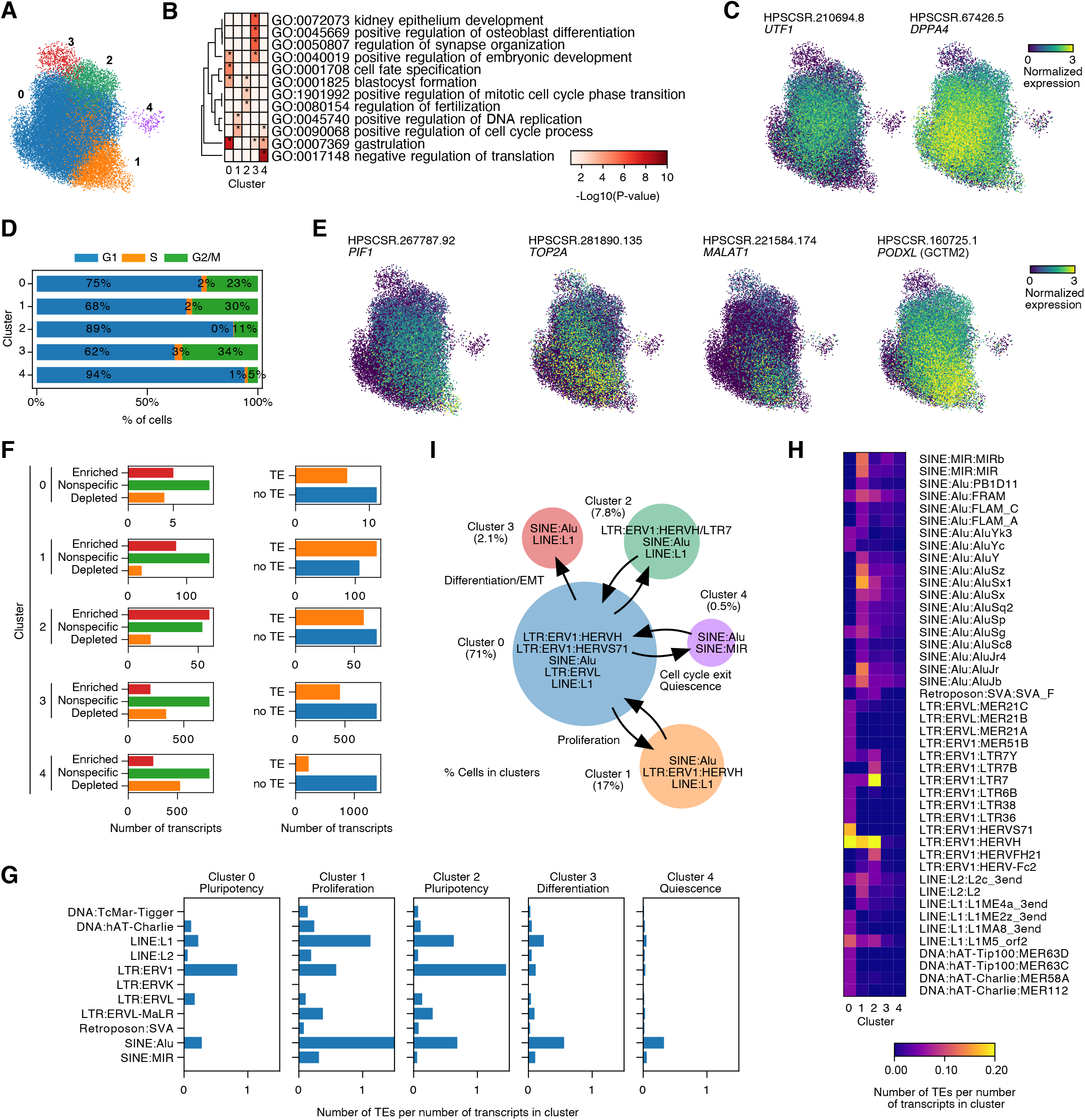
Single cell expression reveals TE containing transcripts are specifically expressed in subpopulations within hPSC cultures. (A) UMAP projection and clustering of scRNA-seq of hPSC data. Clustered using Leiden (resolution=1.0) and clusters showing no differentially regulated genes were iteratively merged. (B) Gene ontology analysis of the genes significantly associated with the indicated clusters (See **Supplemental Table S6** for the list of transcripts). (C) UMAP plots colored by the expression level of two pluripotency genes, *UTF1* and *DPPA4*. (D) Estimates of cell cycle phase of the indicated UMAP clusters based on enrichment of cell cycle genes. (E) UMAP plot colored by expression of the G2/M-associated genes *PIF1*, *TOP2A*, *MALAT1* and *PODXL*. (F) Number of transcripts in each cluster that is enriched, nonspecific or depleted in hPSCs (Left bar charts), and number of transcripts containing a TE (right bar charts). (G) Number of TEs types per number of transcripts in the indicated clusters. (H) Heatmap indicating the number of TE families per number of transcripts in the indicated clusters (I) Schematic indicating the relationship between the hPSC cell sub-populations. The % of cells in each cluster is indicated, and the major TE types expressed are indicated.

Finally, we looked at the composition of TEs and transcripts specific to each of the clusters. Clusters 0, 1 and 2 represent the bulk hPSCs, and exist on a spectrum, from highly proliferating (cluster 1) to medium proliferation (cluster 0) and relatively low proliferation (cluster 2). All three clusters were rich in hPSC-enriched transcripts and TE-containing transcripts (**Fig. 6F**). We measured the frequency of TEs inside the transcripts, and observed a large number of HERVH TEs, and some LINE:L1 and SINE:Alu-containing transcripts specific to cluster 0, 1 and 2 (**Fig. 6G, S8A, B**). Proliferative cluster 1 showed an increase in SINE:Alu containing transcripts (**Fig. 6G**). The two differentiating clusters, 3 and 4, were almost entirely depleted of LTR-containing transcripts (**Fig. 6G**), and were only enriched for SINE:Alu, SINE:MIR and LINE:L1-containing transcripts (**Fig. 6G, H, S8C, D**). Overall, we show that distinct subsets of cells within a hPSC culture contain transcripts with specific patters of TE expression. HERVH TEs are restricted to clusters 0, 1 and 2, and are inversely correlated with proliferation. Proliferating cells tend to express transcripts containing SINEs and LINEs, and cells undergoing an EMT, and starting to differentiate, lose expression of HERVH-containing transcripts (**Fig. 6I**). These results demonstrate the heterogenous patterns of TE expression in single cells in cultures of hPSCs.

## Discussion

TEs constitute the majority of the DNA sequence of the human genome (Hutchins and Pei 2015). They have long been considered as a part, involved in regulatory functions, and providing an evolutionary substrate (Bourque et al. 2018). However, TEs ultimately need to be suppressed and controlled due to their implications in human disease and effects on genome stability (Beck et al. 2010; Hutchins and Pei 2015; Tam et al. 2019). Here, using a combination of short and long read RNA-seq data, we show that TEs are an integral part of the hPSC-transcriptome, contributing to 26% of coding transcripts and 65% of non-coding lncRNA transcripts. The insertion of TEs into coding transcripts is poorly tolerated in the 5’UTR and CDS, but are tolerated or even beneficial for expression inside the 3’UTR. For non-coding transcripts the effect of TEs is uniformly deleterious, and the higher the number of TEs, the lower the level of gene expression.

We found potential coding sequences in our catalog of transcripts; particularly peptides derived from TE insertions. Our estimates of coding transcripts containing a TE suggests that around 20% of these transcripts are capable of generating peptides, at least transiently, although most peptides are not derived directly from the TE sequence. As lincRNAs have been observed to contain potential CDSs (Cabili et al. 2011), it is likely that the non-peptide producing transcripts retain coding signatures but not coding potential. Ultimately, the distinction between coding and non-coding transcripts remains fuzzy, with a large number of genes with unclear designation (Abascal et al. 2018). It is also possible that non-coding transcripts can become coding in a specific cell type. Considering the increasing interest in the biological functions of alternate proteins and small peptides (sCDSs) in other biological contexts (Makarewich and Olson 2017), it will be important to explore how both sCDSs and TE-derived peptides are contributing to hPSCs. One class of TE-containing coding transcript that we could detect peptides for were HERVH-derived viral proteins. Viral proteins have been observed in some cell types, for example HERVW *env* proteins have been found in placental cells, and are thought to help mediate trophoblast invasion into uterine tissues (Frendo et al. 2003; Chuong et al. 2013). hPSCs express HERVH and HERVEs containing RNAs, but we did not observe any peptides. Only HERVK ERVs showed evidence of *gag*, *pol* and *env* translation.

Our catalog of transcripts from hPSCs demonstrates how the complement of TEs is integrated into coding and non-coding transcriptome. Interestingly, the splicing of TEs into pluripotency transcripts has been seen in cancer cells. For example, an AluJb is spliced into the pluripotency gene *LIN28B* and MLT1J into *SALL4* (Yu et al. 2007; Yang et al. 2010; Jang et al. 2019). The splicing of these TEs is thought to convert the pluripotent gene into an oncogene (Jang et al. 2019). Intriguingly, in our hPSC-specific dataset we did not observe these specific TE-transcript splicing patterns (**Supplemental Fig. S9A-C**), suggesting that these are cancer-specific effects, and implying that there is a normal set of TEs that are spliced into transcripts (Fort et al. 2014), and a set that is associated with disease. Indeed, hPSC-specific HERVs were not associated with the expression of pluripotency-related transcripts in human cancers (Zapatka et al. 2020), again suggesting that TE splicing has a normal and diseased pattern. Related to this, the observation of proteins derived from HERVK is noteworthy considering their link with various cancers (Subramanian et al. 2011). The application of long-read sequencing to human diseases, and specifically cancers will be important to shed light on the influence of TEs on transcript splicing in human disease. Overall, our study reveals how the transcribed TE complement of the cell contributes to both coding and non-coding transcriptome of hPSCs, and provides a model of the normal TE transcriptome for comparison to diseased TE expression.

## Authors’ contributions

A.P.H conceived and funded the project, I.A.B., Y.H.L. and A.P.H. performed the bioinformatic analysis. C.M.C., X.L.F., M.G., S.L. and M.D. performed the long-read RNA-seq and validation. B.P.D., Z.L., M.M.A., C.W. performed the scRNA-seq. A.P.H. and I.A.B. wrote the manuscript with assistance from R.J, M.A.E, J.F, J.B.C and C.J. All authors read and approved the manuscript.

## Acknowledgements

This work was supported by the National Natural Science Foundation of China (31970589, 31850410463 to A.P.H., 31801217 to Z.Q., 31850410486 to I.A.B.), the Science and Technology Planning Project of Guangdong Province (2019A050510004 to A.P.H. and R.J.), the National Key Research and Development Program of China (2016YFA0100102 and 2018YFA0106903 to M.A.E.), and the Shenzhen Peacock plan (201701090668B to A.P.H.). Supported by Center for Computational Science and Engineering of Southern University of Science and Technology.

## Accessions

The long-read RNA-seq was deposited in the Sequence Read Archive (SRA) with the accession number: PRJNA631047, and the scRNA-seq data with the accession number: PRJNA631808.

## References

Abascal F, Juan D, Jungreis I, Kellis M, Martinez L, Rigau M, Rodriguez JM, Vazquez J, Tress ML. 2018. Loose ends: almost one in five human genes still have unresolved coding status. Nucleic Acids Res 46: 7070–7084.

Babarinde IA, Li Y, Hutchins AP. 2019. Computational Methods for Mapping, Assembly and Quantification for Coding and Non-coding Transcripts. Comput Struct Biotechnol J 17: 628–637.

Beck CR, Collier P, Macfarlane C, Malig M, Kidd JM, Eichler EE, Badge RM, Moran JV. 2010. LINE-1 retrotransposition activity in human genomes. Cell 141: 1159–1170.

Bourque G, Burns KH, Gehring M, Gorbunova V, Seluanov A, Hammell M, Imbeault M, Izsvak Z, Levin HL, Macfarlan TS et al. 2018. Ten things you should know about transposable elements. Genome Biol 19: 199.

Bulut-Karslioglu A, Macrae TA, Oses-Prieto JA, Covarrubias S, Percharde M, Ku G, Diaz A, McManus MT, Burlingame AL, Ramalho-Santos M. 2018. The Transcriptionally Permissive Chromatin State of Embryonic Stem Cells Is Acutely Tuned to Translational Output. Cell Stem Cell 22: 369–383 e368.

Cabili MN, Trapnell C, Goff L, Koziol M, Tazon-Vega B, Regev A, Rinn JL. 2011. Integrative annotation of human large intergenic noncoding RNAs reveals global properties and specific subclasses. Genes Dev 25: 1915–1927.

Carlevaro-Fita J, Lanzos A, Feuerbach L, Hong C, Mas-Ponte D, Pedersen JS, Drivers P, Functional Interpretation G, Johnson R, Consortium P. 2020. Cancer LncRNA Census reveals evidence for deep functional conservation of long noncoding RNAs in tumorigenesis. Commun Biol 3: 56.

Chen D, Sun N, Hou L, Kim R, Faith J, Aslanyan M, Tao Y, Zheng Y, Fu J, Liu W et al. 2019. Human Primordial Germ Cells Are Specified from Lineage-Primed Progenitors. Cell Rep 29: 4568–4582 e4565.

Chishima T, Iwakiri J, Hamada M. 2018. Identification of Transposable Elements Contributing to Tissue-Specific Expression of Long Non-Coding RNAs. Genes (Basel) 9.

Chuong EB, Rumi MA, Soares MJ, Baker JC. 2013. Endogenous retroviruses function as species-specific enhancer elements in the placenta. Nat Genet 45: 325–329.

Clayton EA, Rishishwar L, Huang TC, Gulati S, Ban D, McDonald JF, Jordan IK. 2020. An atlas of transposable element-derived alternative splicing in cancer. Philos Trans R Soc Lond B Biol Sci 375: 20190342.

Consortium F the RP Clst Forrest AR Kawaji H Rehli M Baillie JK de Hoon MJ Haberle V Lassmann T et al. 2014. A promoter-level mammalian expression atlas. Nature 507: 462–470.

Derrien T, Johnson R, Bussotti G, Tanzer A, Djebali S, Tilgner H, Guernec G, Martin D, Merkel A, Knowles DG et al. 2012. The GENCODE v7 catalog of human long noncoding RNAs: analysis of their gene structure, evolution, and expression. Genome Res 22: 1775–1789.

Dewannieux M, Harper F, Richaud A, Letzelter C, Ribet D, Pierron G, Heidmann T. 2006. Identification of an infectious progenitor for the multiple-copy HERV-K human endogenous retroelements. Genome Res 16: 1548–1556.

Faulkner GJ, Kimura Y, Daub CO, Wani S, Plessy C, Irvine KM, Schroder K, Cloonan N, Steptoe AL, Lassmann T et al. 2009. The regulated retrotransposon transcriptome of mammalian cells. Nat Genet 41: 563–571.

Feng S, Jacobsen SE, Reik W. 2010. Epigenetic reprogramming in plant and animal development. Science 330: 622–627.

Fort A, Hashimoto K, Yamada D, Salimullah M, Keya CA, Saxena A, Bonetti A, Voineagu I, Bertin N, Kratz A et al. 2014. Deep transcriptome profiling of mammalian stem cells supports a regulatory role for retrotransposons in pluripotency maintenance. Nat Genet 46: 558–566.

Frendo JL, Olivier D, Cheynet V, Blond JL, Bouton O, Vidaud M, Rabreau M, Evain-Brion D, Mallet F. 2003. Direct involvement of HERV-W Env glycoprotein in human trophoblast cell fusion and differentiation. Mol Cell Biol 23: 3566–3574.

Fuchs NV, Loewer S, Daley GQ, Izsvak Z, Lower J, Lower R. 2013. Human endogenous retrovirus K (HML-2) RNA and protein expression is a marker for human embryonic and induced pluripotent stem cells. Retrovirology 10: 115.

Goff LA, Rinn JL. 2015. Linking RNA biology to lncRNAs. Genome Res 25: 1456–1465.

Goke J, Lu X, Chan YS, Ng HH, Ly LH, Sachs F, Szczerbinska I. 2015. Dynamic transcription of distinct classes of endogenous retroviral elements marks specific populations of early human embryonic cells. Cell Stem Cell 16: 135–141.

Grow EJ, Flynn RA, Chavez SL, Bayless NL, Wossidlo M, Wesche DJ, Martin L, Ware CB, Blish CA, Chang HY et al. 2015. Intrinsic retroviral reactivation in human preimplantation embryos and pluripotent cells. Nature 522: 221–225.

Hubley R, Finn RD, Clements J, Eddy SR, Jones TA, Bao W, Smit AF, Wheeler TJ. 2016. The Dfam database of repetitive DNA families. Nucleic Acids Res 44: D81–89.

Hutchins AP, Pei D. 2015. Transposable elements at the center of the crossroads between embryogenesis, embryonic stem cells, reprogramming, and long non-coding RNAs. Sci Bull 60: 1722–1733.

Hutchins AP, Yang Z, Li Y, He F, Fu X, Wang X, Li D, Liu K, He J, Wang Y et al. 2017. Models of global gene expression define major domains of cell type and tissue identity. Nucleic Acids Res 45: 2354–2367.

Jang HS, Shah NM, Du AY, Dailey ZZ, Pehrsson EC, Godoy PM, Zhang D, Li D, Xing X, Kim S et al. 2019. Transposable elements drive widespread expression of oncogenes in human cancers. Nat Genet 51: 611–617.

Johnson R, Guigo R. 2014. The RIDL hypothesis: transposable elements as functional domains of long noncoding RNAs. RNA 20: 959–976.

Jonsson ME, Ludvik Brattas P, Gustafsson C, Petri R, Yudovich D, Pircs K, Verschuere S, Madsen S, Hansson J, Larsson J et al. 2019. Activation of neuronal genes via LINE-1 elements upon global DNA demethylation in human neural progenitors. Nat Commun 10: 3182.

Jungreis I, Tress ML, Mudge J, Sisu C, Hunt T, Johnson R, Uszczynska-Ratajczak B, Lagarde J, Wright J, Muir P et al. 2018. Nearly all new protein-coding predictions in the CHESS database are not protein-coding. bioRxiv doi:10.1101/360602: 360602.

Jurka J, Kapitonov VV, Kohany O, Jurka MV. 2007. Repetitive sequences in complex genomes: structure and evolution. Annu Rev Genomics Hum Genet 8: 241–259.

Kammerer U, Germeyer A, Stengel S, Kapp M, Denner J. 2011. Human endogenous retrovirus K (HERV-K) is expressed in villous and extravillous cytotrophoblast cells of the human placenta. J Reprod Immunol 91: 1–8.

Kapusta A, Kronenberg Z, Lynch VJ, Zhuo X, Ramsay L, Bourque G, Yandell M, Feschotte C. 2013. Transposable elements are major contributors to the origin, diversification, and regulation of vertebrate long noncoding RNAs. PLoS Genet 9: e1003470.

Kelley D, Rinn J. 2012. Transposable elements reveal a stem cell-specific class of long noncoding RNAs. Genome Biol 13: R107.

Kilpinen H, Goncalves A, Leha A, Afzal V, Alasoo K, Ashford S, Bala S, Bensaddek D, Casale FP, Culley OJ et al. 2017. Common genetic variation drives molecular heterogeneity in human iPSCs. Nature 546: 370–375.

Kim D, Paggi JM, Park C, Bennett C, Salzberg SL. 2019. Graph-based genome alignment and genotyping with HISAT2 and HISAT-genotype. Nat Biotechnol 37: 907–915.

Kim S, Pevzner PA. 2014. MS-GF+ makes progress towards a universal database search tool for proteomics. Nat Commun 5: 5277.

Kunarso G, Chia NY, Jeyakani J, Hwang C, Lu X, Chan YS, Ng HH, Bourque G. 2010. Transposable elements have rewired the core regulatory network of human embryonic stem cells. Nat Genet 42: 631–634.

Lagarde J, Uszczynska-Ratajczak B, Carbonell S, Perez-Lluch S, Abad A, Davis C, Gingeras TR, Frankish A, Harrow J, Guigo R et al. 2017. High-throughput annotation of full-length long noncoding RNAs with capture long-read sequencing. Nat Genet 49: 1731–1740.

Lau KX, Mason EA, Kie J, De Souza DP, Kloehn J, Tull D, McConville MJ, Keniry A, Beck T, Blewitt ME et al. 2020. Unique properties of a subset of human pluripotent stem cells with high capacity for self-renewal. Nat Commun 11: 2420.

Lev-Maor G, Ram O, Kim E, Sela N, Goren A, Levanon EY, Ast G. 2008. Intronic Alus influence alternative splicing. PLoS Genet 4: e1000204.

Li Q, Hutchins AP, Chen Y, Li S, Shan Y, Liao B, Zheng D, Shi X, Li Y, Chan WY et al. 2017. A sequential EMT-MET mechanism drives the differentiation of human embryonic stem cells towards hepatocytes. Nat Commun 8: 15166.

Lu JY, Shao W, Chang L, Yin Y, Li T, Zhang H, Hong Y, Percharde M, Guo L, Wu Z et al. 2020. Genomic Repeats Categorize Genes with Distinct Functions for Orchestrated Regulation. Cell Rep 30: 3296–3311 e3295.

Ma L, Cao J, Liu L, Du Q, Li Z, Zou D, Bajic VB, Zhang Z. 2019. LncBook: a curated knowledgebase of human long non-coding RNAs. Nucleic Acids Res 47: D128–D134.

Macfarlan TS, Gifford WD, Driscoll S, Lettieri K, Rowe HM, Bonanomi D, Firth A, Singer O, Trono D, Pfaff SL. 2012. Embryonic stem cell potency fluctuates with endogenous retrovirus activity. Nature 487: 57–63.

Makarewich CA, Olson EN. 2017. Mining for Micropeptides. Trends Cell Biol 27: 685–696.

Morillon A, Gautheret D. 2019. Bridging the gap between reference and real transcriptomes. Genome Biol 20: 112.

Naville M, Warren IA, Haftek-Terreau Z, Chalopin D, Brunet F, Levin P, Galiana D, Volff JN. 2016. Not so bad after all: retroviruses and long terminal repeat retrotransposons as a source of new genes in vertebrates. Clin Microbiol Infect 22: 312–323.

Nguyen QH, Lukowski SW, Chiu HS, Senabouth A, Bruxner TJC, Christ AN, Palpant NJ, Powell JE. 2018. Single-cell RNA-seq of human induced pluripotent stem cells reveals cellular heterogeneity and cell state transitions between subpopulations. Genome Res 28: 1053–1066.

Pertea M, Pertea GM, Antonescu CM, Chang TC, Mendell JT, Salzberg SL. 2015. StringTie enables improved reconstruction of a transcriptome from RNA-seq reads. Nat Biotechnol 33: 290–295.

Santoni FA, Guerra J, Luban J. 2012. HERV-H RNA is abundant in human embryonic stem cells and a precise marker for pluripotency. Retrovirology 9: 111.

Schumann GG, Fuchs NV, Tristan-Ramos P, Sebe A, Ivics Z, Heras SR. 2019. The impact of transposable element activity on therapeutically relevant human stem cells. Mob DNA 10: 9.

Steijger T, Abril JF, Engstrom PG, Kokocinski F, Consortium R, Hubbard TJ, Guigo R, Harrow J, Bertone P. 2013. Assessment of transcript reconstruction methods for RNA-seq. Nat Methods 10: 1177–1184.

Subramanian RP, Wildschutte JH, Russo C, Coffin JM. 2011. Identification, characterization, and comparative genomic distribution of the HERV-K (HML-2) group of human endogenous retroviruses. Retrovirology 8: 90.

Tam OH, Ostrow LW, Gale Hammell M. 2019. Diseases of the nERVous system: retrotransposon activity in neurodegenerative disease. Mob DNA 10: 32.

Theunissen TW, Friedli M, He Y, Planet E, O’Neil RC, Markoulaki S, Pontis J, Wang H, Iouranova A, Imbeault M et al. 2016. Molecular Criteria for Defining the Naive Human Pluripotent State. Cell Stem Cell 19: 502–515.

Toker L, Feng M, Pavlidis P. 2016. Whose sample is it anyway? Widespread misannotation of samples in transcriptomics studies. F1000Res 5: 2103.

Wang J, Xie G, Singh M, Ghanbarian AT, Rasko T, Szvetnik A, Cai H, Besser D, Prigione A, Fuchs NV et al. 2014. Primate-specific endogenous retrovirus-driven transcription defines naive-like stem cells. Nature 516: 405–409.

Wang X, You X, Langer JD, Hou J, Rupprecht F, Vlatkovic I, Quedenau C, Tushev G, Epstein I, Schaefke B et al. 2019. Full-length transcriptome reconstruction reveals a large diversity of RNA and protein isoforms in rat hippocampus. Nat Commun 10: 5009.

Wapinski O, Chang HY. 2011. Long noncoding RNAs and human disease. Trends Cell Biol 21: 354–361.

Wheeler TJ, Eddy SR. 2013. nhmmer: DNA homology search with profile HMMs. Bioinformatics 29: 2487–2489.

Wu TD, Watanabe CK. 2005. GMAP: a genomic mapping and alignment program for mRNA and EST sequences. Bioinformatics 21: 1859–1875.

Wucher V, Legeai F, Hedan B, Rizk G, Lagoutte L, Leeb T, Jagannathan V, Cadieu E, David A, Lohi H et al. 2017. FEELnc: a tool for long non-coding RNA annotation and its application to the dog transcriptome. Nucleic Acids Res 45: e57.

Yang J, Gao C, Chai L, Ma Y. 2010. A novel SALL4/OCT4 transcriptional feedback network for pluripotency of embryonic stem cells. PLoS One 5: e10766.

You BH, Yoon SH, Nam JW. 2017. High-confidence coding and noncoding transcriptome maps. Genome Res 27: 1050–1062.

Yu J, Vodyanik MA, Smuga-Otto K, Antosiewicz-Bourget J, Frane JL, Tian S, Nie J, Jonsdottir GA, Ruotti V, Stewart R et al. 2007. Induced pluripotent stem cell lines derived from human somatic cells. Science 318: 1917–1920.

Zapatka M, Borozan I, Brewer DS, Iskar M, Grundhoff A, Alawi M, Desai N, Sultmann H, Moch H, Pathogens P et al. 2020. The landscape of viral associations in human cancers. Nat Genet 52: 320–330.

Zhou T, Benda C, Duzinger S, Huang Y, Li X, Li Y, Guo X, Cao G, Chen S, Hao L et al. 2011. Generation of induced pluripotent stem cells from urine. J Am Soc Nephrol 22: 1221–1228.

