## Supplementary File for "Transposable Element-Gene Splicing Modulates the Transcriptional Landscape of Human Pluripotent Stem Cells"

**Supplemental Table S1 –** Details of the short-read samples used in this study

**Supplemental Table S2** – Primers used for validation

**Supplemental Table S3 –** Details of the short and long-read transcripts assembled in this study

**Supplemental Table S4 –** Summary of the TE content of the assembled transcripts

**Supplemental Table S5** – Peptides matches against the variant coding sequences

**Supplemental Table S6** – Differentially expressed genes in the scRNA-seq clusters

#### Methods

##### Code availability

The full code tree for the analysis presented in this paper can be found at: https://github.com/oaxiom/hesc_lincrna. The code is available under an MIT license. The code repository hosts files that researchers may be interested in:

| | |
|---|---|
| GTF file for the assembly | https://github.com/oaxiom/hesc_lincrna/blob/master/transcript_assembly/gtf/current_gtf.gtf.gz |
| Table of transcripts and TEs | https://github.com/oaxiom/hesc_lincrna/blob/master/te_discovery/te_transcripts/transcript_table_merged.mapped.tsv.gz |
| Transcript feature table | https://github.com/oaxiom/hesc_lincrna/blob/master/transcript_assembly/feature_table/assembly_hPSC_detailed.tsv.gz |
| Peptide mass spec results | https://github.com/oaxiom/hesc_lincrna/blob/master/massspec/results_gene.tsv.gz |

##### Cell lines, cell culture methods and experimental techniques

The cell lines used in this study were hESCs (H1 and WIBR3 line) and iPSCs (c11/S0730 line; (Zhou et al. 2011)). These cell lines were grown under typical growth conditions in mTeSR1 (Stemcell technologies: 85850) on pre-coated Matrigel plates (Corning: 354277). Medium was replaced every 24 hours. Cells were passaged by single-cell Accutase (SIGMA: A6964) every 5 days.

**RNA extraction, long read RNA-seq and RT-PCR**

Total RNA was extracted using Trizol (MRC: RN190). The concentration of the extracted RNA was measured by Nano-drop. For long read RNA-seq, we first confirmed the cell status by RT-PCR of marker genes such as Sox2 and Nanog. We then sent the samples for sequencing. Long read sequencing for the two cell lines were done in duplicates. For qPCR of selected transcripts, 1 ug of extracted RNA was retro-transcribed using the PrimeScript RT Master Mix (Taraka: RR036A). Thereafter, cDNA samples were amplified by real-time PCR (qRT-PCR) using TB Green™ Premix Ex Taq™ II (Taraka: RR820A) with the primers listed in **Supplemental Table S2**.

**Source of the RNA-seq and deepCAGE data sets**

The RNA-seq data were obtained from two sources. The short reads were obtained from sequence read archives (SRA). Out of 317 publicly available short read datasets of wild-type unperturbed hESC and iPSC samples, 150 could pass the stringent quality tests. First, we discarded samples with single ended reads. We then aligned the remaining 197 paired-end samples with HISAT2 (v2.1.0) (Kim et al. 2019). As a measure of read quality, we retained 171 samples that have at least 70% alignment rate. To ascertain cell identity, we ran global gene expression correlation. We also checked the expression of the stem cell specific genes *NANOG* and *SOX2* in addition to *GATA4, SOX17, and SOX1,* which mark differentiation. These procedures led to discard an additional 21 samples, leaving 150 good quality hPSC samples. We also sequenced long reads in duplicate from hESCs (H1) and iPSCs (c11/S0730 line; (Zhou et al. 2011)) using the PacBio sequencing platform. The number of reads for each sample are presented in **Supplemental Table S1.** Three replicates each of the alignments of deepCAGE data of both hESC and iPSC were downloaded from FANTOM5 project (Abugessaisa et al. 2017).

**Pipeline for transcript assembly**

Transcripts were assembled independently from SR and LR, and then combined (**Supplemental Fig. S1E**). For SR, the samples of hESC and iPSC were first filtered to ensure sample reliability. Then the reads were mapped to human genome hg38 using HISAT2 (Kim et al. 2019). The alignments were then merged with SAMtools (Li et al. 2009). StringTie (Pertea et al. 2015) was used to assemble the transcripts from the merged alignment using GENCODE v32 annotations. Transcripts with no inferred strand were discarded. For each LR sample, consensus sequences were first generated from subread data using *ccs* (Gordon et al. 2015). Next, *lima* was used to generate full length reads by primer removal and demultiplexing. Next, noise from full length reads was removed using isoseq3. The noiseless alignment files were converted to FASTA format using BAMtools (Barnett et al. 2011). Noise-free full-length reads were then aligned to the human genome hg38 assembly using GMAP (Wu and Watanabe 2005). The alignments from the four samples were merged using SAMTools. StringTie was then used to assemble the transcript from the alignments. As with SR, any transcript that could not be assigned to a strand was discarded. We then merged the SR and LR transcripts. The expression levels of the merged transcripts were then computed from merged and individual SR alignments. The expression level and the spread of expression among the 150 SR samples were further used to filter the set of transcripts.

#Preparing reference for HISAT2

```
$ extract_splice_sites.py Homo_sapiens.GRCh38.91.gtf >Homo_sapiens.GRCh38.91.ss
$ extract_exons.py Homo_sapiens.GRCh38.91.gtf >Homo_sapiens.GRCh38.91.exon
$ hisat2-build --ss Homo_sapiens.GRCh38.91.ss --exon Homo_sapiens.GRCh38.91.exon
Homo_sapiens.GRCh38.dna.primary_assembly.fa hisat2_hg38
```

### Hisat2 alignment for each sample

```
$ hisat2 -p 6 --dta -x hisat2_hg38 -1 SRR597894_1.fq.gz -2 SRR597894_2.fq.gz -S
SRR597894.sam
$ samtools sort -@ 6 -o SRR597894.bam SRR597894.sam
```

#Transcript assembly with SR

```
$ samtools merge -@ 32 -nurlf –h SRR597894.bam -b good150bam_list stem_merge150good.bam
$ samtools sort -@ 32 -o stem_merge150good_sorted.bam stem_merge150good.bam
$ stringtie stem_merge150good_sorted.bam -G gencode.v32.annotation.gtf -o short_read_150_samples.gtf -p 32 -c 1 -v &> short_read_150_samples.out
```

### Isoseq pipeline for LR samples

```
$ ccs --numThreads 8 --noPolish --minPasses 1 H1_m54299_181207_142926.subreads.bam H1_m54299_181207_142926.subreads.ccs_np.bam

$ lima H1_m54299_181207_142926.subreads.ccs_np.bam m54296_181207_094757.adapters.3p.fasta H1_m54299_181207_142926.subreads.fl.bam --guess --isoseq --no-pbi --num-threads 32

$ isoseq3 refine H1_m54299_181207_142926.subreads.ccs_np.bam m54296_181207_094757.adapters.5p.fasta H1_m54299_181207_142926.subreads.flnc.bam

$ bamtools convert -format fasta –in H1_m54299_181207_142926_reqA.subreads.flnc.bam -out H1_m54299_181207_142926_reqA.subreads.flnc.fasta
```

#Alignment for LR samples, using GMAP

```
$ gmap_build -k 8 -d ens_hg38_vs91_gmap Homo_sapiens.GRCh38.dna.primary_assembly.fa
$ gmap -D ens_hg38_vs91_gmap -t 6 -d ens_hg38_vs91_gmap -f samse --max-intronlength-middle 2000000 --localsplicedist 4000000 --totallength 4000000 H1_m54299_181207_142926_reqA.subreads.flnc.fasta > H1_m54299_181207_142926_reqA.subreads.flnc.bam
$samtools merge -@ 32 -nurlf -h merged26.sam -b LR_bam_list LR_merged.bam
$samtools sort -@ 32 -o LR_merged_sorted.bam LR_merged.bam
```

#Transcript assembly for LR samples

```
$ stringtie LR_merged_sorted.bam -G gencode.v32.annotation.gtf –o long_read_4_samples.gtf -p 32 -c 1 -v &> long_read_4_samples.out
```

#Comparing SR and LR gtf files, done after unstranded transcripts were discarded

```
$ python2 compare_gtf.py –i short_read_150_samples_stranded.gtf -r
long_read_4_samples_stranded.gtf -o SR_LR_matched.all -m SR_LR_matched.paired
```

#Merge SR and LR gtf files, using the result of compare_gtf.py

```
$ python2 update_sr_lr.py –s short_read_150_samples_stranded.gtf –l
long_read_4_samples_stranded.gtf –m SR_LR_matched.paired –o SR_LR_merged.gtf
```

#Transcript quantifications for merged bam and individual SR samples

```
$ stringtie stem_merge150good_sorted.bam -G SR_LR_updated_200bp.gtf -o
SR_LR_updated_200bp_count.gtf -e -B
$ stringtie SRR597894.bam -G SR_LR_200bp.gtf -o SR_LR_SRR597894.gtf -e -B
```

#Extracting expression matrix. The input is a directory of all samples from Stringtie quantification

```
$ python2 expression_count_all.py -i count_dir -c TPM -o SR_LR_updated_200bp_TPM_count.tsv
```

**Transcript coding potential**

The coding potential of the transcripts were assessed with FEELnc (Wucher et al. 2017). The transcripts of protein-coding and lincRNA biotypes of GENCODE transcripts were used as training data set for FEELnc. Using the training dataset, FEELnc decided the coding potential threshold of 0.432 (**Supplemental Fig. S2G**) for protein-coding transcripts.

### FEELnc coding potential

```
$ FEELnc_codpot.pl -i assembly_hPSC_final.gtf -a gencode.v32.pc_transcripts.fa -l
gencode.v32.lncRNA_transcripts.fa -g Homo_sapiens.GRCh38.dna.primary_assembly.fa  --
outname=assembly_hPSC_final –outdir=FEELnc_use
```

**DeepCAGE data analyses**

The three biological replicates of each of hESC and iPSC samples (Consortium et al. 2014) were merged and indexed using SAMtools. The bam files were then converted to wiggle files using deeptools bamCoverage (Ramirez et al. 2014). Then the wiggle file and the transcript files were used to compute the matrix using deeptools computeMatrix. The matrices were plotted using deeptools plotHeatmap. For the read coverage, regions covering 100bp upstream and downstream of the transcription start sites (TSS) of hPSC transcripts were extracted. The estimation of the number of reads mapped to each region in the two merged alignments was done using deeptools multiBamSummary.

**Bodymap data analyses**

The sequence read data for the 16 body map samples were downloaded and aligned by HISAT2 (Derrien et al. 2012). The expression levels (TPM) were computed for each sample, using the same StringTie quantification that was used for hPSCs. The standard deviation of log2-transformed expression TPM levels for each transcript was then computed. The z-score was then computed for the expression level of each hPSC transcript using the mean and the standard deviation computed from bodymap samples. Transcripts with z-score < -1 are classified as "depleted"; those with z-score > 1 are classified as "enriched", while those with -1≤ z-score ≤1 are classified as "nonspecific".

**Detection of TE insertions inside transcripts**

For each transcript, the FASTA sequence was extracted for the assembled transcript. An edited version of dfam (Hubley et al. 2016) database of TE HMM models was used, with all non-primate TE families removed. Then, nhmmer was used to search against this transcript database;

```
$ nhmmer -E 1e-10 --cpu 32 --dna --noali --tblout tblout.$out.tsv Dfam_edited.hmm $inp
>/dev/null
```

As nhmmer can discover overlapping TEs, these were trimmed based on a logical progression of criteria: (1) If two domains entirely contained each other, then the TE with the lowest E-value was kept. (2) If the percentage of overlap between two pairs of TEs was > 60%, then the domain with the best E-value was retained and the other TE deleted. This was repeated iteratively across all pairs of domains until both conditions were satisfied. The final Supplemental Table of TE-containing transcripts is in **Supplemental Table S3**, and at https://github.com/oaxiom/hesc_lincrna/blob/master/te_discovery/te_transcripts/transcript_table_merged.mapped.tsv.gz.

**Determination of coding sequences inside novel coding transcripts and detection of peptides from TE-derived putative coding sequences in LC-MS/MS data**

FEELnc only provides an overall coding potential score, and does not obtain a location of the probable coding sequence. To estimate the likely location of the coding sequence we took the longest open reading frame from any ATG to a STOP codon. Note that for a coding frame to be guessed, it must contain an ATG, although we would add in-frame STOP codons if the transcript appeared to be truncated (this is to match the transcript annotations in the GENCODE transcripts, which also contain STOPs up to the end of the transcript). We then compared the CDS calls against the CDSs reported in the GENCODE transcripts, and 85% of our GENCODE calls matched the CDS reported by GENCODE. Finally, as a final filter to remove any novel CDSs that actually matched to a full-length GENCODE CDS, the resulting proteins were blasted against the GENCODE protein FASTA, and any matches >95% that matched the full length of the protein were deleted from the analysis. This final filter sounds like it should not be necessary, but our transcript assembly contains transcripts with no STOP and the GENCODE assembly contains truncated transcripts that may have both no STOP nor ATG codon. Consequently, simply matching up locations and sizes of the coding sequences would not report these truncated protein sequences.

We then took the set of proteins that contained a TE, had a valid CDS, and did not have a BLAST hit covering >95% identity for the full-length protein. We looked at sub-hits within the putative coding region, and masked out (using the '-' character) any regions of the protein that had a >95% identity with any fragment of a protein inside the GENCODE protein FASTA. We further deleted any proteins with <20 amino acids, and any proteins that did not contain at least one K, L or R amino acid that could be cleaved by the Lys-C/trypsin digestion combination used in the HipSci proteomics data (Kilpinen et al. 2017) (PRIDE accession: PXD003903).

The HipSci mzml data was converted to centroid (peakPicker) files using the MSGF+ (Kim and Pevzner 2014) command:

```
$ msconvert $in --filter "peakPicking cwt msLevel=1-" --mzML --32 --outfile $out.cwt.mzML
```

Using the final set of proteins and the HipSci data we then performed a search of the LC-MS/MS spectra using MSGF+ (Kim and Pevzner 2014) with the command:

```
$ MSGFPlus.jar –s $in.cwt.mzML –thread 30 –d masked_peptides.fa –e 3 –inst 1 –t 20ppm –ti
1,2 –mod mods.txt –ntt 2 –tda 1 –o out.mzid
```

To reduce the search space and reduce false positive hits, the modifications (mods.txt) considered included only Carbamidomethyl and Oxidation. A peptide was considered a hit if the q-value reported by MSGF+ was <0.05. Peptide hits for transcripts are in **Supplemental Table S5**.

**Single cell RNA-seq and analysis**

We performed sc-RNA-seq of hPSC samples using 10x genomics platform, according to the manufacturer's instructions. We also added the WTC iPSC scRNA-seq data from E-MTAB-6687 (Nguyen et al. 2018), and two UCLA1 hESC line samples from GSE140021 (Chen et al. 2019). Both studies had used the 10x single cell platform. As 10x-based sc-RNA-seq is heavily biased to the 3' ends of transcripts, we reduced the set of total transcripts to only their unique 3' ends (within 200 bp either side of the TTS). Where transcripts shared an overlapping 3'end one of the ends was kept. This reduced the number of transcripts to 88,520 (87%) of the total set of transcripts. The sc-RNA-seq reads were aligned to the hg38 genome using STARsolo (Dobin et al. 2013), with the settings and using the appropriate whitelist barcode file (version 1 for (Nguyen et al. 2018), and version 2 for (Chen et al. 2019) and our data):

```
$ STAR --runRNGseed 42 --runThreadN 12 --readFilesCommand zcat  --outFilterMultimapNmax 100 --winAnchorMultimapNmax 100 --outSAMmultNmax 1 --outSAMtype BAM SortedByCoordinate --twopassMode Basic --soloType Droplet --soloFeatures Gene --soloBarcodeReadLength 0  --soloCBwhitelist versionX.txt --outSAMattributes NH HI AS nM CR CY UR UY --genomeDir gtf_index/SAindex --outFileNamePrefix ss.$out --readFilesIn $p1 $p2
```

The BAM was then processed using te_counter (https://github.com/oaxiom/te_counter), which is a lightweight reimplementation of the scTE algorithm (He et al., manuscript in preparation):

### Make index, only needs to be done once:

```
$ te_genome -m custom -g 3ends_only.idx --gtf ends.gtf
```

### Count the reads from the BAM file, demultiplexing the barcodes and UMIs. $white is the whitelist of barcodes from the 10x platform and must be correct for the version of 10x used.

```
$ te_count -m custom -g 3ends_only.idx -i $inp -o $out.tsv --sc --se --strand -w $white
```

We then used SCANPY (Wolf et al. 2018) to process the data. Cells with less than 1500 genes or 3000 counts, and those with more than 50,000 or 8500 genes were deleted. Transcripts expressed in less than 100 cells were excluded. The resulting data was normalized using SCRAN (Lun et al. 2016). UMAPs were generated based on the first 20 principal components, and Leiden clustering was performed. Differential expression was called using the rank_genes_groups SCANPY function, with the settings: "method='t-test_overestim_var', n_genes=10000". Significantly different genes were kept if their FDR corrected q-value was <0.05 and they were >2-fold in any cluster.

#### References


Abugessaisa I, Noguchi S, Hasegawa A, Harshbarger J, Kondo A, Lizio M, Severin J, Carninci P, Kawaji H, Kasukawa T. 2017. FANTOM5 CAGE profiles of human and mouse reprocessed for GRCh38 and GRCm38 genome assemblies. *Sci Data* **4**: 170107.

Barnett DW, Garrison EK, Quinlan AR, Stromberg MP, Marth GT. 2011. BamTools: a C++ API and toolkit for analyzing and managing BAM files. *Bioinformatics* **27**: 1691-1692.

Chen D, Sun N, Hou L, Kim R, Faith J, Aslanyan M, Tao Y, Zheng Y, Fu J, Liu W et al. 2019. Human Primordial Germ Cells Are Specified from Lineage-Primed Progenitors. *Cell Rep* **29**: 4568-4582 e4565.

Consortium F the RP Clst Forrest AR Kawaji H Rehli M Baillie JK de Hoon MJ Haberle V Lassmann T et al. 2014. A promoter-level mammalian expression atlas. *Nature* **507**: 462-470.

Derrien T, Johnson R, Bussotti G, Tanzer A, Djebali S, Tilgner H, Guernec G, Martin D, Merkel A, Knowles DG et al. 2012. The GENCODE v7 catalog of human long noncoding RNAs: analysis of their gene structure, evolution, and expression. *Genome Res* **22**: 1775-1789.

Dewannieux M, Harper F, Richaud A, Letzelter C, Ribet D, Pierron G, Heidmann T. 2006. Identification of an infectious progenitor for the multiple-copy HERV-K human endogenous retroelements. *Genome Res* **16**: 1548-1556.

Dobin A, Davis CA, Schlesinger F, Drenkow J, Zaleski C, Jha S, Batut P, Chaisson M, Gingeras TR. 2013. STAR: ultrafast universal RNA-seq aligner. *Bioinformatics* **29**: 15-21.

Gordon SP, Tseng E, Salamov A, Zhang J, Meng X, Zhao Z, Kang D, Underwood J, Grigoriev IV, Figueroa M et al. 2015. Widespread Polycistronic Transcripts in Fungi Revealed by Single-Molecule mRNA Sequencing. *PLoS One* **10**: e0132628.

Hubley R, Finn RD, Clements J, Eddy SR, Jones TA, Bao W, Smit AF, Wheeler TJ. 2016. The Dfam database of repetitive DNA families. *Nucleic Acids Res* **44**: D81-89.

Jang HS, Shah NM, Du AY, Dailey ZZ, Pehrsson EC, Godoy PM, Zhang D, Li D, Xing X, Kim S et al. 2019. Transposable elements drive widespread expression of oncogenes in human cancers. *Nat Genet* **51**: 611-617.

Kilpinen H, Goncalves A, Leha A, Afzal V, Alasoo K, Ashford S, Bala S, Bensaddek D, Casale FP, Culley OJ et al. 2017. Common genetic variation drives molecular heterogeneity in human iPSCs. *Nature* **546**: 370-375.

Kim D, Paggi JM, Park C, Bennett C, Salzberg SL. 2019. Graph-based genome alignment and genotyping with HISAT2 and HISAT-genotype. *Nat Biotechnol* **37**: 907-915.

Kim S, Pevzner PA. 2014. MS-GF+ makes progress towards a universal database search tool for proteomics. *Nat Commun* **5**: 5277.

Li H, Handsaker B, Wysoker A, Fennell T, Ruan J, Homer N, Marth G, Abecasis G, Durbin R, Genome Project Data Processing S. 2009. The Sequence Alignment/Map format and SAMtools. *Bioinformatics* **25**: 2078-2079.

Lun AT, McCarthy DJ, Marioni JC. 2016. A step-by-step workflow for low-level analysis of single-cell RNA-seq data with Bioconductor. *F1000Res* **5**: 2122.

Nguyen QH, Lukowski SW, Chiu HS, Senabouth A, Bruxner TJC, Christ AN, Palpant NJ, Powell JE. 2018. Single-cell RNA-seq of human induced pluripotent stem cells reveals cellular heterogeneity and cell state transitions between subpopulations. *Genome Res* **28**: 1053-1066.

Pertea M, Pertea GM, Antonescu CM, Chang TC, Mendell JT, Salzberg SL. 2015. StringTie enables improved reconstruction of a transcriptome from RNA-seq reads. *Nat Biotechnol* **33**: 290-295.

Ramirez F, Dundar F, Diehl S, Gruning BA, Manke T. 2014. deepTools: a flexible platform for exploring deep-sequencing data. *Nucleic Acids Res* **42**: W187-191.

Wolf FA, Angerer P, Theis FJ. 2018. SCANPY: large-scale single-cell gene expression data analysis. *Genome Biol* **19**: 15.

Wu TD, Watanabe CK. 2005. GMAP: a genomic mapping and alignment program for mRNA and EST sequences. *Bioinformatics* **21**: 1859-1875.

Wucher V, Legeai F, Hedan B, Rizk G, Lagoutte L, Leeb T, Jagannathan V, Cadieu E, David A, Lohi H et al. 2017. FEELnc: a tool for long non-coding RNA annotation and its application to the dog transcriptome. *Nucleic Acids Res* **45**: e57.

Zhou T, Benda C, Duzinger S, Huang Y, Li X, Li Y, Guo X, Cao G, Chen S, Hao L et al. 2011. Generation of induced pluripotent stem cells from urine. *J Am Soc Nephrol* **22**: 1221-1228.

**Supplemental Figure S1 – Pipeline for the assembly of high quality hPSC transcripts**

(A) Ranked alignment rate for all short-read samples. The line indicates our cutoff of >70% of reads must map to the hg38 human genome assembly. 171 samples passed, 26 failed.

(B) Co-correlation Pearson $R^2$ heatmap for all passing samples

(C) Example plot showing all short-read samples and the expression of pluripotency genes (*SOX2*, *NANOG*) and differentiation genes (*SOX17*, *SOX1* and *GATA4*). Samples were removed if they expressed high levels of differentiated marker genes.

(D) Co-correlation Pearson $R^2$ heatmap for final set of 150 short-read RNA-seq samples.

(E) Pipeline used for hPSC transcript assembly

(F) Transcript filtering stages and results for getting high-quality hPSC transcripts

(G) Genome view of example long-read transcripts that span an intron, and are not supported by short-read data, and likely represent unspliced transcripts.

(H) Percent of LR and SR- based transcripts that passed the quality control filters.

**Supplemental Figure S2 – Validation of the transcript assembly and non-coding RNA determination.**

(A) Gel electrophoresis result for 40 selected hPSC transcripts

(B) DeepCAGE data confirms TSS of many transcripts. DeepCAGE data was from (Consortium et al. 2014; Abugessaisa et al. 2017).

(C) Scatter plot indicating the Relationship between RNA-seq expression levels and deepCAGE tag counts

(D) Completeness (percent of exons or splices of the closest GENCODE transcript that were correctly retrieved), of exon and splice concordance versus GENCODE.

(E) Exon and splice completeness for transcripts with different read support.

(F) Percentage of transcripts in each class with SR, LR or SR+LR support.

(G) FEELnc coding probability plot for determination of coding potential threshold

(H) Exon and splice percentage concordance versus GENCODE for matching and variant transcripts for coding and non-coding transcripts.

(I) Expression spread, the percentage of SR samples with expression, for coding and non-coding transcripts

(J) Number of transcripts that are enriched, nonspecific or depleted in hPSC versus bodymap expression

(K) Distribution of hPSC transcript class based on bodymap expression

**Supplemental Figure S3 – Frequency of splicing of TEs into coding genes.**

(A) Relative density of selected TE subtypes in transcripts' 5' UTR, CDS or 3'UTR. Each plot is normalized to the length of the transcript region. The blue line indicates all GENCODE transcripts, the red, green and yellow lines indicate hPSC-specific expression categories.

(B) As in panel b, but for specific families of TEs.

(C) Domain plots for selected transcripts containing a L1HS_5end. The location of TEs in the LINE, SINE LTR or DNA/SVAs (Retroposons) are indicated. The CDS is indicted by a think black box (unless it is non-coding), and the numbers below the 'coding seq' indicate the start of the indicated regions in mRNA coordinates. Locations of splice sites are indicated by a slanted red line.

(D) As in panel c, but showing LTR7 and HERVH TEs

(E) As in panel c, but showing LTR7 and HERVH in the middle of transcripts.

**Supplemental Figure S4 – Mass spec peptides derived from HERVK transcripts**

(A) Domain plots for selected transcripts containing a HERVH or LTR7. The location of TEs in the LINE, SINE LTR or DNA/SVAs are indicated. The CDS is indicted by a think black box, and the numbers below the 'coding seq' indicate the start of the indicated regions in mRNA coordinates. Locations of splice sites are indicated by a slanted red line. The vertical red boxes indicate the positions for MS-detected peptides.

(B) Alignment of the mass spec detected peptides (marked in red) against a hypothetical progenitor version of the HERVK viral proteins *gag*, *pol* and *env* (Dewannieux et al. 2006).

**Supplemental Figure S5 – PCAT14 is the origin of many HERVK-derived peptides**

(A) Genome view of the PCAT14 transcripts detected in hPSCS (top rows), along with the short-read RNA-seq pileup data (middle row, blue), the location of TEs (red bars = LTRs, green=SINEs). The lower rows are the GENCODE transcripts.

(B) CLUSTAL-omega alignment of *gag*, and *pol* HERVK viral proteins against putative ORFs from *PCAT14*.

**Supplemental Figure S6 – Density of TE insertions across lncRNAs.**

(A) Frequency of TEs subtypes across the normalized length of all lncRNAs. TSS=Transcription start site; TTS = Transcription termination site. Frequency is normalized to the total number of transcripts in each expression class.

(B) As in panel A, but for LINE families.

(C) As in panel A, but for SINE families.

(D) As in panel A, but for Unknown and Satellite families.

(E) As in panel A, but for the DNA TE Tigger1.

(F) As in panel A, but for ERVK and ERVL-MaLR TEs.

(G) As in panel A, but for ERV1 LTRs.

**Supplemental Fig. S7 – Analysis of the scRNA-seq data.**

(A) Pileup of sc-RNA-seq data across the 3' TTS ends of the transcripts. The pileups indicate the signal for + stranded transcripts on both the + and – strand (left pileups), and the same for the – strandred transcripts (right-hand pileups). The pileups are centered on the TTS and extend 2 kb in either direction. The lower heatmaps show the same data as the pileups, but each row is a transcript.

(B) UMAP plots showing the distributions of hESCs and iPSCs in the sc-RNA-seq data.

(C) As in panel B, but each cell is colored by the sample it originated from.

(D) UMAP plots colored by gene expression intensity. Pluripotency genes are shown

(E) As in panel D, but for epithelial genes (*CDH1*, *EPCAM*)

(F) As in panel D, but for *CD9*, a marker for proliferation.

(G) As in panel D, but for mesenchymal genes (*CDH2*, VIM*)*.

**Supplemental Fig. S8 – LTR:ERV1:HERVH-containing transcripts are associated with high proliferating hPSCs**.

(A) UMAP plots (left) colored by transcript expression for a cluster 0/2 gene (HPSCLR.4844.3). The domain plot is shown on the right-hand side, it indicates the location of the inserted TEs in the transcript. The 'coding seq' part indicates the presence or absence of a CDS (this transcript is predicted to be non-coding).

(B) As in panel B, but for cluster 1-specific transcripts.

(C) As in panel B, but for a cluster 4-specific transcript.

(D) As in panel B, but for a cluster 3-specific transcript. The tick part of the 'coding seq' indicates the CDS.

**Supplemental Fig. S9 – TE-gene chimera transcripts of the pluripotency genes *SALL4* and *LIN28B*, expressed in cancer cells, cannot be assembled in hPSCs**.

(A) Genome views containing the transcripts assembled in this study (top), read pileups for the sc-RNA-seq (red and orange pileups), the short-read RNA-seq data (blue), and the locations of LINEs, SINEs, LTRs and Retroposons in the genome. The GENCODE transcripts are shown below. The grey box indicates the position of the MLT1J spliced into *SALL4* as detected in Ref.(Jang et al. 2019).

(B) As in panel a, but showing the *LIN28A* locus. The location of the Alu's spliced into *LIN28A* detected in Ref. (Jang et al. 2019) is indicated.

(C) All of the domain maps for *SALL4* detected in this study. The location of TEs in the LINEs, SINEs, LTRs or DNA/SVAs are indicated. The CDS is indicted by a think black box, and the numbers below the 'coding seq' indicate the start of the indicated regions in mRNA coordinates. Locations of splice sites are indicated by a slanted red line.
