## Supplementary Figures for "Transposable Element-Gene Splicing Modulates the Transcriptional Landscape of Human Pluripotent Stem Cells"

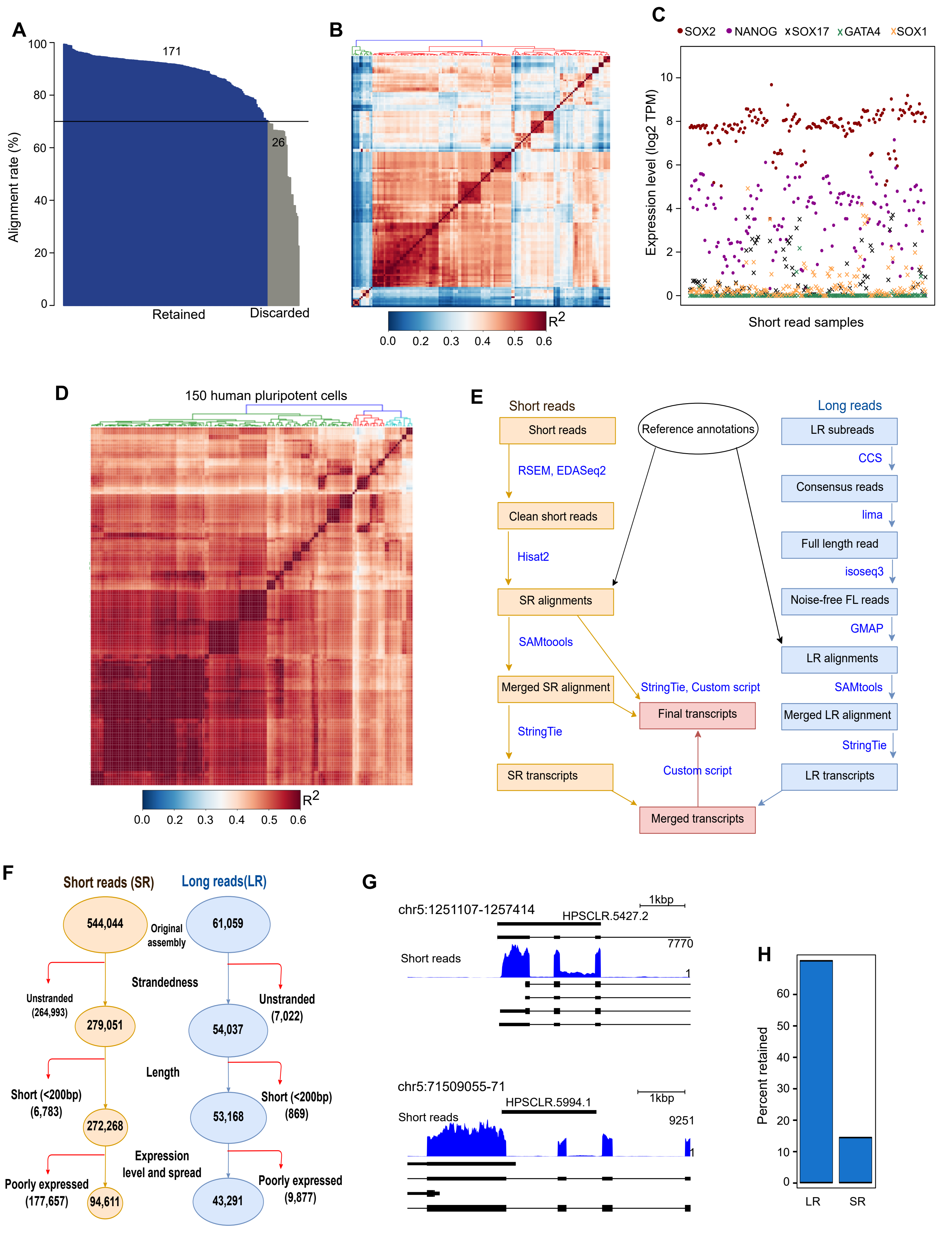
A
Alignment rate (%)
171
26
Retained
Discarded
B
$R^2$
C
SOX2
NANOG
SOX17
GATA4
SOX1
Expression level (log2 TPM)
Short read samples
D
150 human pluripotent cells
$R^2$
E
Short reads
Short reads
Reference annotations
Long reads
LR subreads
CCS
Consensus reads
RSEM, EDASeq2
Clean short reads
lima
Full length read
Hisat2
isoseq3
SR alignments
Noise-free FL reads
GMAP
SAMtoools
LR alignments
Merged SR alignment
StringTie, Custom script
SAMtools
Final transcripts
Merged LR alignment
StringTie
StringTie
Custom script
SR transcripts
LR transcripts
Merged transcripts
F
Short reads (SR)
Long reads(LR)
544,044
Original assembly
61,059
Strandedness
Unstranded (264,993)
Unstranded (7,022)
279,051
54,037
Length
Short (<200bp) (6,783)
Short (<200bp) (869)
272,268
53,168
Expression level and spread
Poorly expressed (177,657)
Poorly expressed (9,877)
94,611
43,291
G
chr5:1251107-1257414
1kbp
HPSCLR.5427.2
7770
Short reads
chr5:71509055-71
1kbp
HPSCLR.5994.1
Short reads
9251
H
Percent retained
LR
SR

**Figure S1**

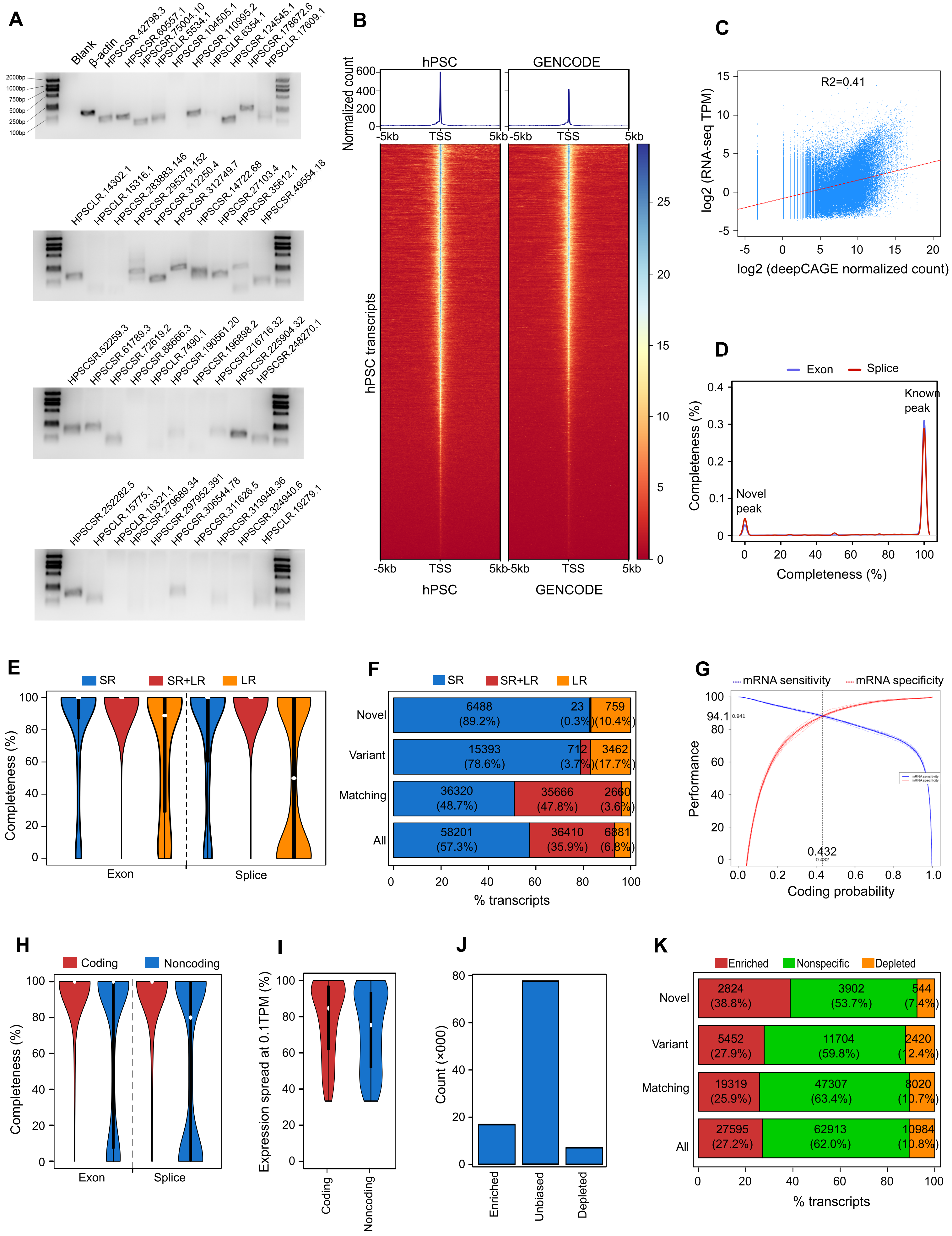

A
B
C
D
E
F
G
H
I
J
K
hPSC
GENCODE
Normalized count
hPSC transcripts
TSS
R2=0.41
log2 (RNA-seq TPM)
log2 (deepCAGE normalized count)
Exon
Splice
Known peak
Novel peak
Completeness (%)
SR
SR+LR
LR
Novel
Variant
Matching
All
% transcripts
mRNA sensitivity
mRNA specificity
Performance
Coding probability
Coding
Noncoding
Expression spread at 0.1TPM (%)
Count (×000)
Enriched
Unbiased
Depleted
Nonspecific


**Figure S2**

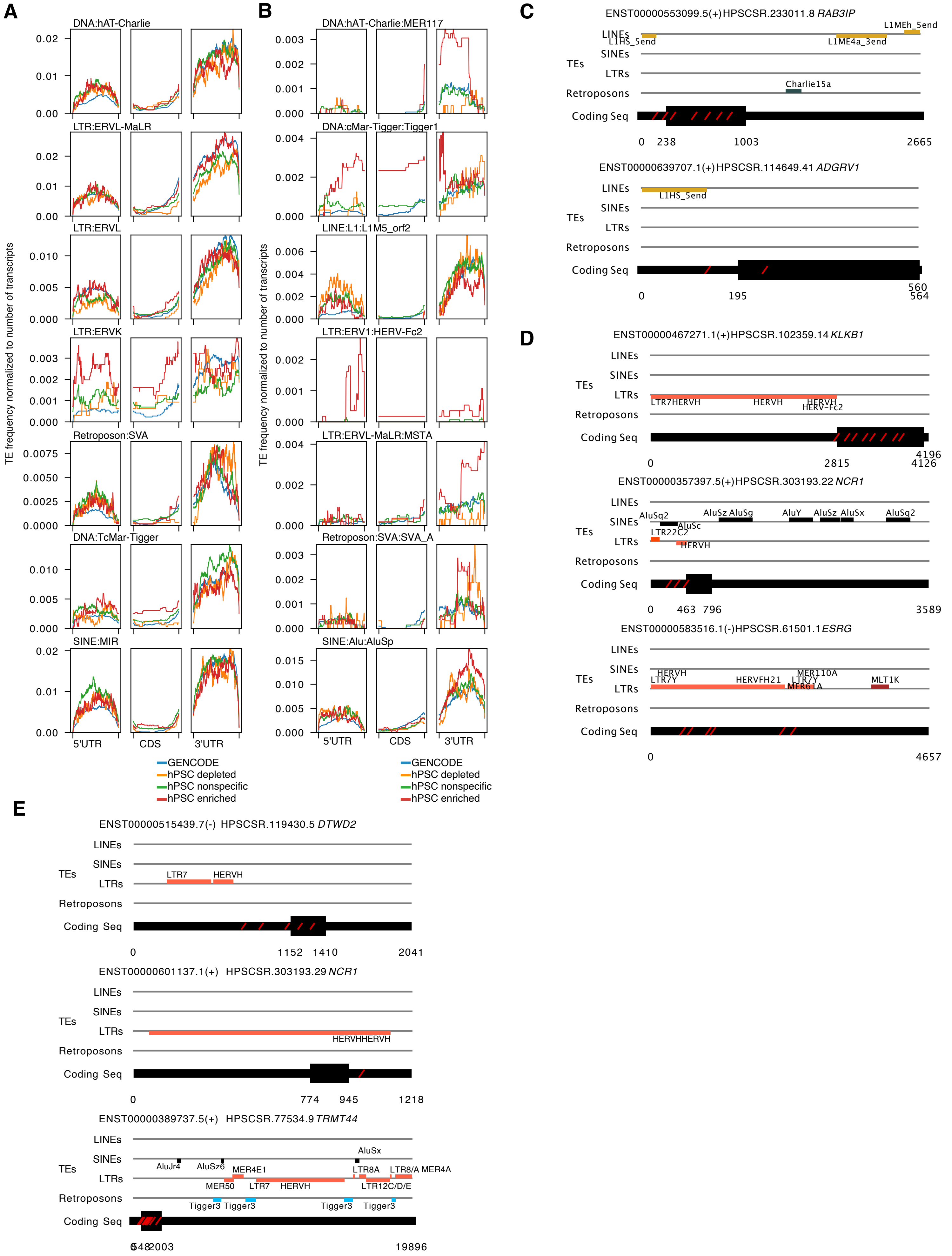
A
B
C
D
E
DNA:hAT-Charlie
LTR:ERVL-MaLR
LTR:ERVL
LTR:ERVK
Retroposon:SVA
DNA:TcMar-Tigger
SINE:MIR
DNA:hAT-Charlie:MER117
DNA:cMar-Tigger:Tigger1
LINE:L1:L1M5_orf2
LTR:ERV1:HERV-Fc2
LTR:ERVL-MaLR:MSTA
Retroposon:SVA:SVA_A
SINE:Alu:AluSp
TE frequency normalized to number of transcripts
5'UTR
CDS
3'UTR
GENCODE
hPSC depleted
hPSC nonspecific
hPSC enriched
ENST00000553099.5(+)HPSCSR.233011.8 RAB3IP
ENST00000639707.1(+)HPSCSR.114649.41 ADGRV1
ENST00000467271.1(+)HPSCSR.102359.14 KLKB1
ENST00000357397.5(+)HPSCSR.303193.22 NCR1
ENST00000583516.1(-)HPSCSR.61501.1 ESRG
ENST00000515439.7(-) HPSCSR.119430.5 DTWD2
ENST00000601137.1(+) HPSCSR.303193.29 NCR1
ENST00000389737.5(+) HPSCSR.77534.9 TRMT44
TEs
LINEs
SINEs
LTRs
Retroposons
Coding Seq

**Figure S3**

**A**

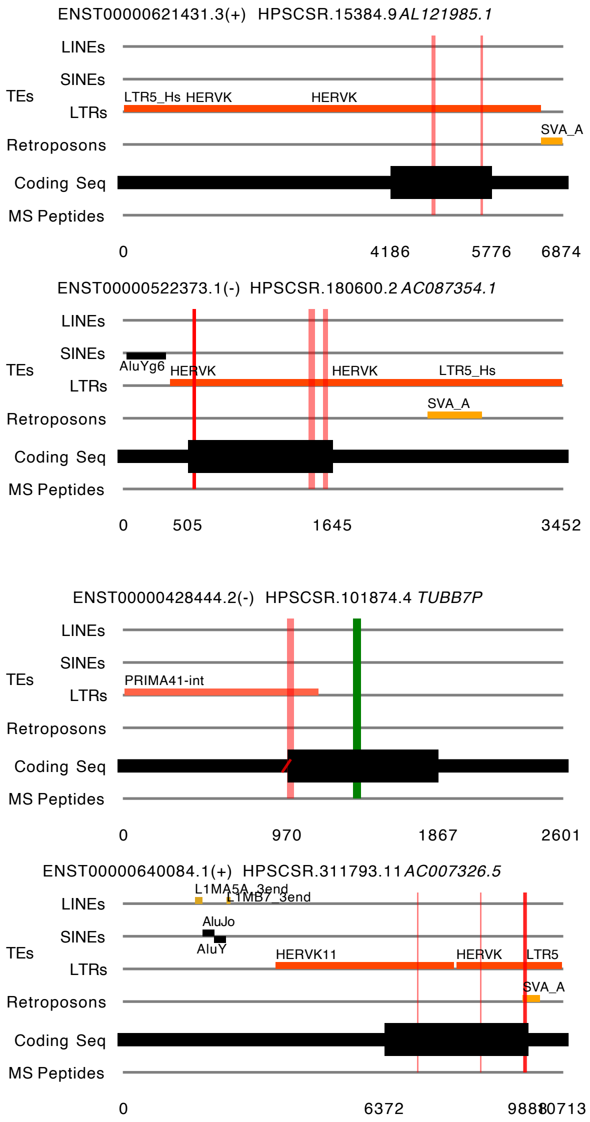


**B**

*gag* (17 unique peptides)

```
            YASYLSFIK
MGQTKSKIKSKYASYLSFIKILLKRGGVKVSTKNLIKLFQIIEQFCPWFPEQGTLDLKDW

                                                 SCVTDCEEEAG
                             ATLEPFQTGEDIVSVSDAPK
               IIPLTVWNDWAIIK
KRIGKELKQAGRKGNIIPLTVWNDWAIIKAALEPFQTEEDSVSVSDAPGSCIIDCNENTR

TESQQGTESSHCK                             LGEGGPESLGPSEPK
KKSQKETEGLHCEYVAEPVMAQSTQNVDYNQLQEVIYPETLKLEGKGPELVGPSESKPRG

TQPLVVYQYR
TSPLPAGQVPVTLQPQKQVKENKTQPPVAYQYWPPAELQYRPPPESQYGYPGMPPAPQGR

APYPQPPTRRLNPTAPPSRQGSELHEIIDKSRKEGDTEAWQFPVTLEPMPPGEGAQEGEP

                                            LTPYDWEILAK
                                            LIPYDWEILAK
                                 TLLDSIAHGNR           SSLSP
                DMKEGVKQYGPNSPYMR                      SSLSS
PTVEARYKSFSIKMLKDMKEGVKQYGPNSPYMRTLLDSIAHGHRLIPYDWEILAKSSLSP


SQYLQFK
SQFLQFK
       TWWIDGVQEQVR
SQFLQFKTWWIDGVQEQVRRNRAANPPVNIDADQLLGIGQNWSTISQQALMQNEAIEQVR

         IQDPGTAFPINSI-R    EPYPDFVAR
AICLRAWEKIQDPGSTCPSFNTVRQGSKEPYPDFVARLQDVAQKSIADEKARKVIVELMA

                  VPAGVDVITEYVK
YENANPECQSAIKPLKGKVPAGSDVISEYVKACDGIGGAMHKAMLMAQAITGVVLGGQVR

TFGGKCYNCGQIGHLKKNCPVLNKQNITIQATTTGREPPDLCPRCKKGKHWASQCRSKFD

           RGQPQAPQQTGAFPVQLFVPQGFQGQQPLQK
KNGQPLSGNEQRGQPQAPQQTGAFPIQPFVPQGFQGQQPPLSQVFQGISQLPQYNNCPPP

QAAVQQ
```

*pol* (2 unique peptides)

```
NKSKKRRNRVSFLGAATVEPPKPIPLTWKTEKPVWVNQWPLPKQKLEALHLLANEQLEKG

HIEPSFSPWNSPVFVIQKKSGKWRMLTDLRAVNAVIQPMGPLQPGLPSPAMIPKDWPLII

IDLKDCFFTIPLAEQDCEKFAFTIPAINNKEPATRFQWKVLPQGMLNSPTICQTFVGRAL

QPVREKFSDCYIIHYIDDILCAAETKDKLIDCYTFLQAEVANAGLAIASDKIQTSTPFHY

                                                         GDS
LGMQIENRKIKPQKIEIRKDTLKTLNDFQKLLGDINWIRPTLGIPTYAMSNLFSILRGDS

DLNSKRMLTPEATKEIK
DLNSKRMLTPEATKEIKLVEEKIQSAQINRIDPLAPLQLLIFATAHSPTGIIIQNTDLVE

WSFLPHSTVKTFTLYLDQIATLIGQTRLRIIKLCGNDPDKIVVPLTKEQVRQAFINSGAW

QIGLANFVGIIDNHYPKTKIFQFLKLTTWILPKITRREPLENALTVFTDGSSNGKAAYTG

PKERVIKTPYQSAQRAELVAVITVLQDFDQPINIISDSAYVVQATRDVETALIKYSMDDQ

              RNFPFYITHIR
LNQLFNLLQQTVRKRNFPFYITHIRAHTNLPGPLTKANEQADLLVSSALIKAQELHALTH

VNAAGLKNKFDVTWKQAKDIVQHCTQCQVLHLPTQEAGVNPRGLCPNALWQMDVTHVPSF

GRLSYVHVTVDTYSHFIWATCQTGESTSHVKKHLLSCFAVMGVPEKIKTDNGPGYCSKAF

QKFLSQWKISHTTGIPYNSQGQAIVERTNRTLKTQLVKQKEGGDSKECTTPQMQLNLALY

TLNFLNIYRNQTTTSAEQHLTGKKNSPHEGKLIWWKDNKNKTWEIGKVITWGRGFACVSP

GENQLPVWIPTRHLKFYNEPIRDAKKSTSAETETPQSSTVDSQDEQNGDVRRTDEVAIHQ

EGRAADLGTTKEADAVSYKISREHKGDTNPREYAACSLDDCINGGKSPYACRSSCS
```

*env* (4 unique peptides)

```
MNPSEMQRKAPPR             MNKMVTSEEQMKLPSTK
MNPSEMQRKAPPRRRRHRNRAPLTHKMNKMVTSEEQMKLPSTKKAEPPTWAQLKKLTQLA

TKYLENTKVTQTPESMLLAALMIVSMVVSLPMPAGAAAANYTYWAYVPFPPLIRAVTWMD

NPIEVYVNDSVWVPGPIDDRCPAKPEEEGMMINISIGYRYPPICLGRAPGCLMPAVQNWL

VEVPTVSPISRFTYHMVSGMSLRPRVNYLQDFSYQRSLKFRPKGKPCPKEIPKESKNTEV

LVWEECVANSAVILQNNEFGTIIDWAPRGQFYHNCSGQTQSCPSAQVSPAVDSDLTESLD

LQSFYPWEWGEK                             LMMAQSPIRIWK
KHKHKKLQSFYPWEWGEKGISTPRPKIISPVSGPEHPELWRLTVASHHIRIWSGNQTLET

RDRKPFYTVDLNSSLTVPLQSCVKPPYMLVVGNIVIKPDSQTITCENCRLLTCIDSTFNW

QHRILLVRAREGVWIPVSMDRPWEASPSIHILTEVLKGVLNRSKRFIFTLIAVIMGLIAV

TATAAVAGVALHSSVQSVNFVNDWQKNSTRLWNSQSSIDQKLANQINDLRQTVIWMGDRL

MSLEHRFQLQCDWNTSDFCITPQIYNESEHHWDMVRRHLQGREDNLTLDISKLKEQIFEA

SKAHLNLVPGTEAIAGVADGLANLNPVTWVKTIGSTTIINLILILVCLFCLLLVCRCTQQ

LRRDSDHRERAMMTMAVLSKRKGGNVGKSKRDQIVTVSV
```

**Figure S4**

**A**

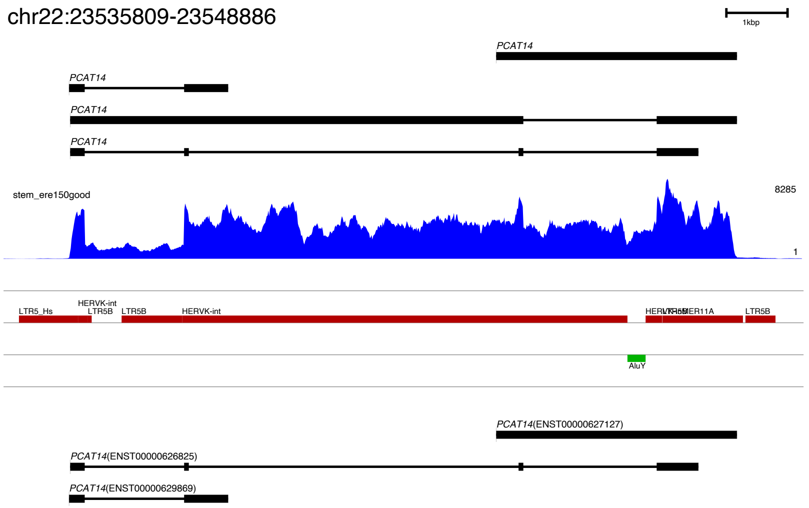


**B**

```
PCAT14      ----MGQTESKYASYLSFIKILLRRGGVRASTENLITLFQTIEQFCPWFPEQGTLDLKDW      56
GAG         MGQTKSKIKSKYASYLSFIKILLKRGGVKVSTKNLIKLFQIIEQFCPWFPEQGTLDLKDW      60
                 .: :*************:****:.**:***.*** ******************

PCAT14      EKIGKELKQANREGKIIPLTVWNDWAIIKATLEPFQTGEDIVSVSDAPKSCVTDCEEEAG      116
GAG         KRIGKELKQAGRKGNIIPLTVWNDWAIIKAALEPFQTEEDSVSVSDAPGSCIIDCNENTR      120
            ::********.*:*:***************:****** ** ******* **: **:*::

PCAT14      TESQQGTESSHCKYVAESVMAQSTQNVDYSQLQEIIYPESSKLGEGGPESLGPSEPKPRS      176
GAG         KKSQKETEGLHCEYVAEPVMAQSTQNVDYNQLQEVIYPETLKLEGKGPELVGPSESKPRG      180
            .:**: **. **:**** ***********.****:****: **   *** :**** ***.

PCAT14      PSTPPPVVQMPVTLQPQTQVRQAQTPRENQVERDRVSIPAMPTQIQYPQYQPVENKTQPL      236
GAG         TS-PLPAGQVPVTLQPQKQV--------------------------------KENKTQPP      207
             * * *. *:*******.**                                ******

PCAT14      VVYQYRLPTELQYRPPSEVQYRPQAVCPVPNSTAPYQQPTAMASNSPATQDAALYPQPPT      296
GAG         VAYQYWPPAELQYRPPPESQYGYPGMPPAP-------------------QGRAPYPQPPT      248
            *.***  *:******* * **   .: *.*                   *. * ******

PCAT14      VRLNPTASRSGQGGALHAVIDEARKQGDLEAWRFLVILQLVQAGEETQVGAPARAETRCE      356
GAG         RRLNPTAPPSRQGSELHEIIDKSRKEGDTEAWQFPVTLEPMPPGEGAQEGEPPTVEARYK      308
             ******  * **. ** :**::**:** ***:* * *: :   ** :* * *  .*:* :

PCAT14      PFTMKMLKDIKEGVKQYGSNSPYIRTLLDSIAHGNRLTPYDWEILAKSSLSSSQYLQFKT      416
GAG         SFSIKMLKDMKEGVKQYGPNSPYMRTLLDSIAHGHRLIPYDWEILAKSSLSPSQFLQFKT      368
             *::*****:******** ****:**********:** ************* **:*****

PCAT14      WWIDGVQEQVRKNQATKPTVNIDADQLLGTGPNWSTINQQSVMQNEAIEQVRAICLRAWG      476
GAG         WWIDGVQEQVRRNRAANPPVNIDADQLLGIGQNWSTISQQALMQNEAIEQVRAICLRAWE      428
            ***********:*:*::* ********** * *****.**::*****************

PCAT14      KIQDPGTAFP-INSIRQGSKEPYPDFVARLQDAAQKSITDDNARKVIVELMAYENANPEC      535
GAG         KIQDPGSTCPSFNTVRQGSKEPYPDFVARLQDVAQKSIADEKARKVIVELMAYENANPEC      488
            ******:: * :*::*****************.*****:*::*****************

PCAT14      QSAIKPLKGKVPAGVDVITEYVKACDGIGGAMHKAMLMAQAMRGLTLGGQVRTFGKKCYN      595
GAG         QSAIKPLKGKVPAGSDVISEYVKACDGIGGAMHKAMLMAQAITGVVLGGQVRTFGGKCYN      548
            ************** ***:*********************: *:.********* ****

PCAT14      CGQIGHLKRSCPGLNKQNIINQAINSKK--------------------------------      623
GAG         CGQIGHLKKNCPVLNKQNITIQATTTGREPPDLCPRCKKGKHWASQCRSKFDKNGQPLSG      608
            ********:.** ******  ** .: :

PCAT14      ----------------------------------------------------------      623
GAG         NEQRGQPQAPQQTGAFPIQPFVPQGFQGQQPPLSQVFQGISQLPQYNNCPPPQAAVQQ      666
```

**C**

```
PCAT14_orf-57_POL      ------------------------------------------------------------ 0
PCAT14_orf-91_POL      ------------------------------------------------------------ 0
PCAT14_orf-52_POL      ------------------------------------------------------------ 0
POL                    NKSKKRRNRVSFLGAATVEPPKPIPLTWKTEKPVWVNQWPLPKQKLEALHLLANEQLEKG 60

PCAT14_orf-57_POL      ------------------------------------------------------------ 0
PCAT14_orf-91_POL      ------------------------------------------------------------ 0
PCAT14_orf-52_POL      ------------------------------------------------------------ 0
POL                    HIEPSFSPWNSPVFVIQKKSGKWRMLTDLRAVNAVIQPMGPLQPGLPSPAMIPKDWPLII 120

PCAT14_orf-57_POL      ------------------------------------------------------------ 0
PCAT14_orf-91_POL      ------------------------------------------------------------ 0
PCAT14_orf-52_POL      ------------------------------------------------------------ 0
POL                    IDLKDCFFTIPLAEQDCEKFAFTIPAINNKEPATRFQWKVLPQGMLNSPTICQTFVGRAL 180

PCAT14_orf-57_POL      ------------------------------------------------------------ 0
PCAT14_orf-91_POL      ------------------------------------------------------------ 0
PCAT14_orf-52_POL      ------------------------------------------------------------ 0
POL                    QPVREKFSDCYIIHYIDDILCAAETKDKLIDCYTFLQAEVANAGLAIASDKIQTSTPFHY 240

PCAT14_orf-57_POL      ------------------------------------------------------------ 0
PCAT14_orf-91_POL      ------------------------------------------------------------ 0
PCAT14_orf-52_POL      ------------------------------------------------MSNLFSFLRGDS 12
POL                    LGMQIENRKIKPQKIEIRKDTLKTLNDFQKLLGDINWIRPTLGIPTYAMSNLFSILRGDS 300

PCAT14_orf-57_POL      ------------------------------------------------------------ 0
PCAT14_orf-91_POL      ------------------------------------------------------------ 0
PCAT14_orf-52_POL      ELNSERTLTPEATKEIKLIEEKIRSAQVNRIDHLAPLQILIFATAHSLTGIIVQNTDLVE 72
POL                    DLNSKRMLTPEATKEIKLVEEKIQSAQINRIDPLAPLQLLIFATAHSPTGIIIQNTDLVE 360

PCAT14_orf-57_POL      ------------------------------------------------------------ 0
PCAT14_orf-91_POL      ------------------------------------------------------------ 0
PCAT14_orf-52_POL      WSFLPHSTIKTFTLYLDQMATLIGQGRL-------------------------------- 100
POL                    WSFLPHSTVKTFTLYLDQIATLIGQTRLRIIKLCGNDPDKIVVPLTKEQVRQAFINSGAW 420

PCAT14_orf-57_POL      ------------------------------------------------------------ 0
PCAT14_orf-91_POL      ------------------------------------------------------------ 0
PCAT14_orf-52_POL      ------------------------------------------------------------ 100
POL                    QIGLANFVGIIDNHYPKTKIFQFLKLTTWILPKITRREPLENALTVFTDGSSNGKAAYTG 480

PCAT14_orf-57_POL      ------------------------------------------------------------ 0
PCAT14_orf-91_POL      --------------------------------------------------------MDDQ 4
PCAT14_orf-52_POL      ------------------------------------------------------------ 100
POL                    PKERVIKTPYQSAQRAELVAVITVLQDFDQPINIISDSAYVVQATRDVETALIKYSMDDQ 540

PCAT14_orf-57_POL      ------------------------------------------------------------ 0
PCAT14_orf-91_POL      LNPLFNLLQQNVRKRNFPFYITHIRAHTNLPGPLTKANEQADLLVSSAFMEAQELHALTH 64
PCAT14_orf-52_POL      ------------------------------------------------------------ 100
POL                    LNQLFNLLQQTVRKRNFPFYITHIRAHTNLPGPLTKANEQADLLVSSALIKAQELHALTH 600

PCAT14_orf-57_POL      ------------------------------------------------------------ 0
PCAT14_orf-91_POL      VNAIGLKNKFDITWKQTKNIVQHCTQCQILHLATQEARVNPRGLCPNVLWQMDVMHVPSF 124
PCAT14_orf-52_POL      ------------------------------------------------------------ 100
POL                    VNAAGLKNKFDVTWKQAKDIVQHCTQCQVLHLPTQEAGVNPRGLCPNALWQMDVTHVPSF 660

PCAT14_orf-57_POL      ------------------------------------------------------------ 0
PCAT14_orf-91_POL      GKLSFVHVTVDTYSHFIWATCQTGESTSHVKRHLLSCFPVMGVPEKVKTDNGPGYCSKAV 184
PCAT14_orf-52_POL      ------------------------------------------------------------ 100
POL                    GRLSYVHVTVDTYSHFIWATCQTGESTSHVKKHLLSCFAVMGVPEKIKTDNGPGYCSKAF 720

PCAT14_orf-57_POL      ----------------------------------------------------MQLNLALY 8
PCAT14_orf-91_POL      QKFLNQWKITHTIGILYNSQGQAIIERTNRTLKAQLVKQKKGKDRSI-TLPRCNLI---- 239
PCAT14_orf-52_POL      ------------------------------------------------------------ 100
POL                    QKFLSQWKISHTTGIPYNSQGQAIVERTNRTLKTQLVKQKEGGDSKECTTPQMQLNLALY 780

PCAT14_orf-57_POL      TLNVLNIYRNQTTTSAEQHLTGKRNSPHEGKLIWWKDNKNKTWEMGKVITWGRGFACVSP 68
PCAT14_orf-91_POL      ------------------------------------------------------------ 239
PCAT14_orf-52_POL      ------------------------------------------------------------ 100
POL                    TLNFLNIYRNQTTTSAEQHLTGKKNSPHEGKLIWWKDNKNKTWEIGKVITWGRGFACVSP 840

PCAT14_orf-57_POL      GENQLPVWIPTRHLKFYNELTGDAKKSVEMET--PQSTRQST-----GVSSSSKETATSK 121
PCAT14_orf-91_POL      ------------------------------------------------------------ 239
PCAT14_orf-52_POL      ------------------------------------------------------------ 100
POL                    GENQLPVWIPTRHLKFYNEPIRDAKKSTSAETETPQSSTVDSQDEQNGDVRRTDEVAIHQ 900

PCAT14_orf-57_POL      NGP-----------------------------------------------------     124
PCAT14_orf-91_POL      --------------------------------------------------------     239
PCAT14_orf-52_POL      --------------------------------------------------------     100
POL                    EGRAADLGTTKEADAVSYKISREHKGDTNPREYAACSLDDCINGGKSPYACRSSCS     956
```

### Figure S5

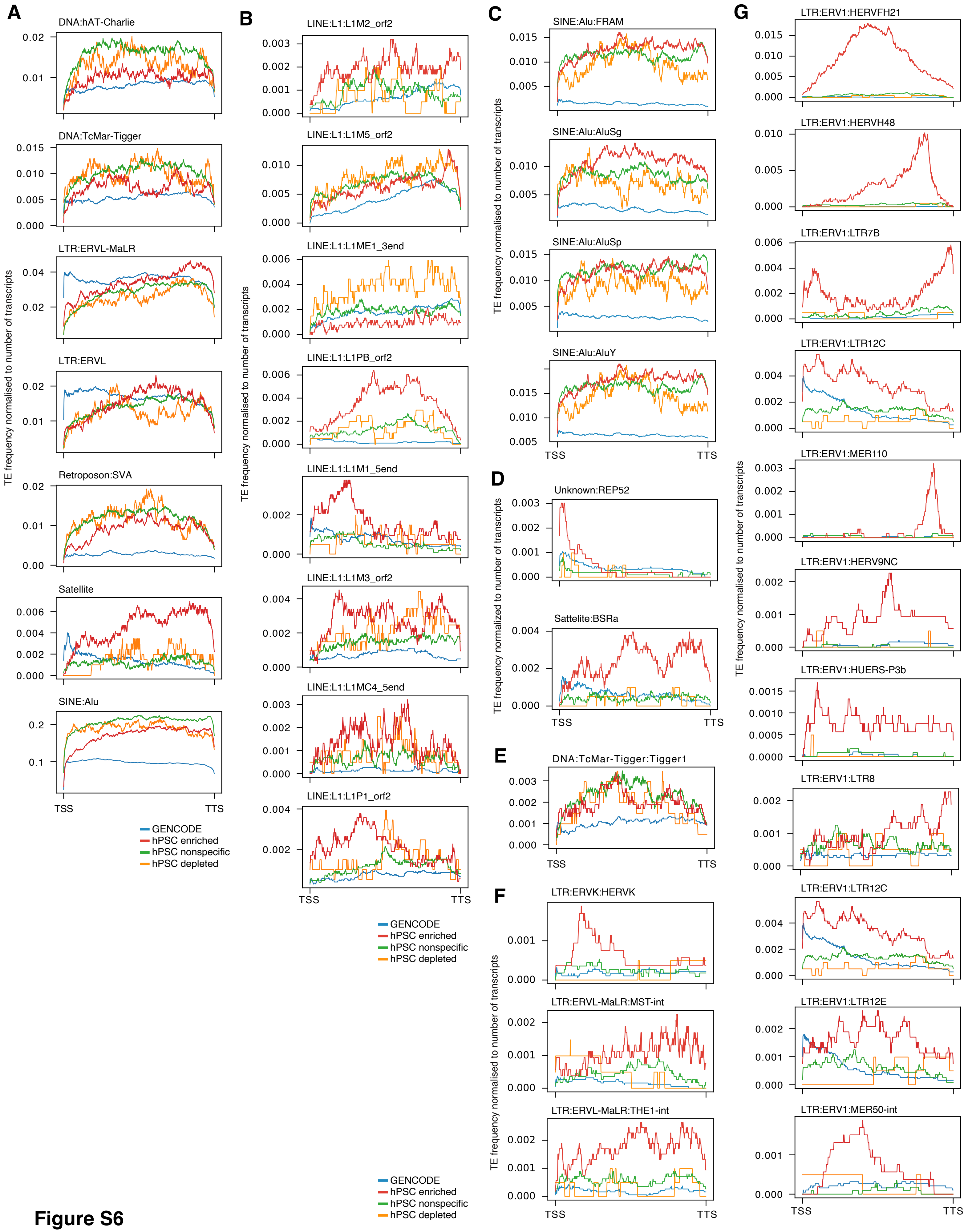
A
DNA:hAT-Charlie
DNA:TcMar-Tigger
LTR:ERVL-MaLR
LTR:ERVL
Retroposon:SVA
Satellite
SINE:Alu
TE frequency normalised to number of transcripts
TSS
TTS
GENCODE
hPSC enriched
hPSC nonspecific
hPSC depleted
B
LINE:L1:L1M2_orf2
LINE:L1:L1M5_orf2
LINE:L1:L1ME1_3end
LINE:L1:L1PB_orf2
LINE:L1:L1M1_5end
LINE:L1:L1M3_orf2
LINE:L1:L1MC4_5end
LINE:L1:L1P1_orf2
TE frequency normalised to number of transcripts
TSS
TTS
GENCODE
hPSC enriched
hPSC nonspecific
hPSC depleted
C
SINE:Alu:FRAM
SINE:Alu:AluSg
SINE:Alu:AluSp
SINE:Alu:AluY
TE frequency normalised to number of transcripts
TSS
TTS
D
Unknown:REP52
Sattelite:BSRa
TE frequency normalized to number of transcripts
TSS
TTS
E
DNA:TcMar-Tigger:Tigger1
TSS
TTS
F
LTR:ERVK:HERVK
LTR:ERVL-MaLR:MST-int
LTR:ERVL-MaLR:THE1-int
TE frequency normalised to number of transcripts
TSS
TTS
GENCODE
hPSC enriched
hPSC nonspecific
hPSC depleted
G
LTR:ERV1:HERVFH21
LTR:ERV1:HERVH48
LTR:ERV1:LTR7B
LTR:ERV1:LTR12C
LTR:ERV1:MER110
LTR:ERV1:HERV9NC
LTR:ERV1:HUERS-P3b
LTR:ERV1:LTR8
LTR:ERV1:LTR12C
LTR:ERV1:LTR12E
LTR:ERV1:MER50-int
TE frequency normalised to number of transcripts
TSS
TTS

**Figure S6**

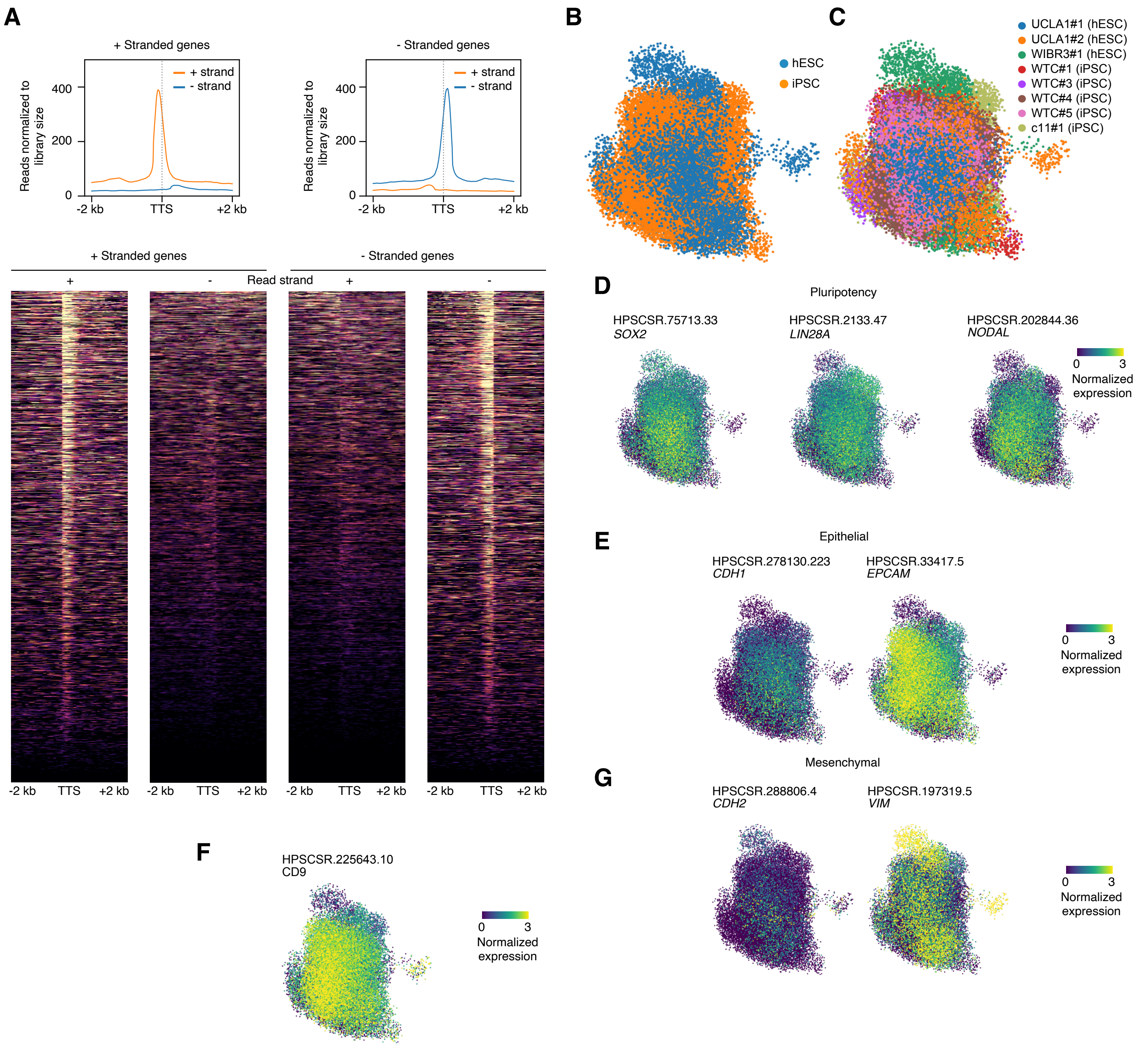
A
+ Stranded genes
- Stranded genes
Reads normalized to library size
400
200
0
+ strand
- strand
-2 kb
TTS
+2 kb
Read strand
+
-
B
hESC
iPSC
C
UCLA1#1 (hESC)
UCLA1#2 (hESC)
WIBR3#1 (hESC)
WTC#1 (iPSC)
WTC#3 (iPSC)
WTC#4 (iPSC)
WTC#5 (iPSC)
c11#1 (iPSC)
D
Pluripotency
HPSCSR.75713.33
SOX2
HPSCSR.2133.47
LIN28A
HPSCSR.202844.36
NODAL
0
3
Normalized expression
E
Epithelial
HPSCSR.278130.223
CDH1
HPSCSR.33417.5
EPCAM
G
Mesenchymal
HPSCSR.288806.4
CDH2
HPSCSR.197319.5
VIM
F
HPSCSR.225643.10
CD9

**Figure S7**

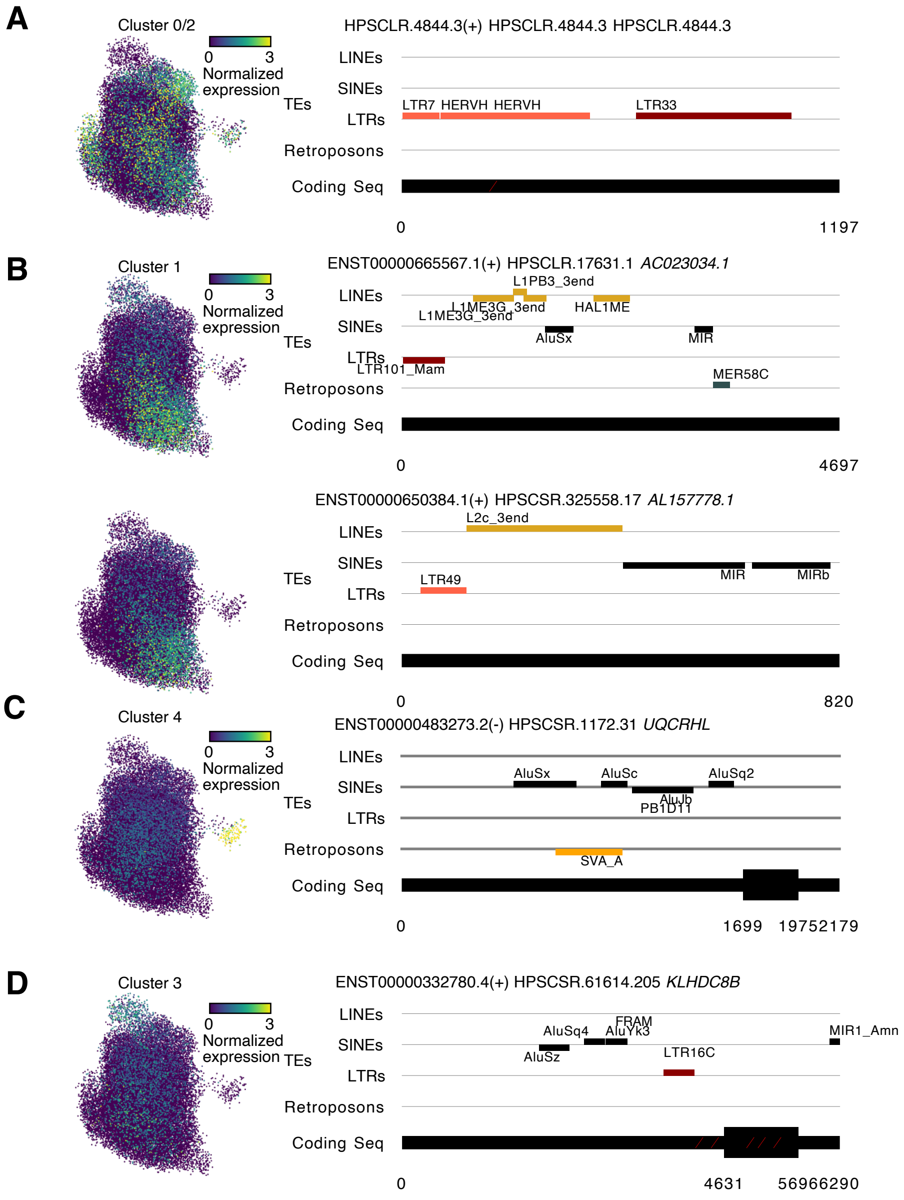
A
Cluster 0/2
0 3 Normalized expression
HPSCLR.4844.3(+) HPSCLR.4844.3 HPSCLR.4844.3
LINEs
SINEs
TEs
LTRs
LTR7 HERVH HERVH
LTR33
Retroposons
Coding Seq
0
1197
B
Cluster 1
ENST00000665567.1(+) HPSCLR.17631.1 AC023034.1
L1PB3_3end
L1ME3G_3end
HAL1ME
AluSx
MIR
LTR101_Mam
MER58C
4697
ENST00000650384.1(+) HPSCSR.325558.17 AL157778.1
L2c_3end
MIR
MIRb
LTR49
820
C
Cluster 4
ENST00000483273.2(-) HPSCSR.1172.31 UQCRHL
AluSx
AluSc
AluSq2
AluJb
PB1D11
SVA_A
1699
19752179
D
Cluster 3
ENST00000332780.4(+) HPSCSR.61614.205 KLHDC8B
AluSq4
FRAM
AluYk3
AluSz
MIR1_Amn
LTR16C
4631
56966290

**Figure S8**

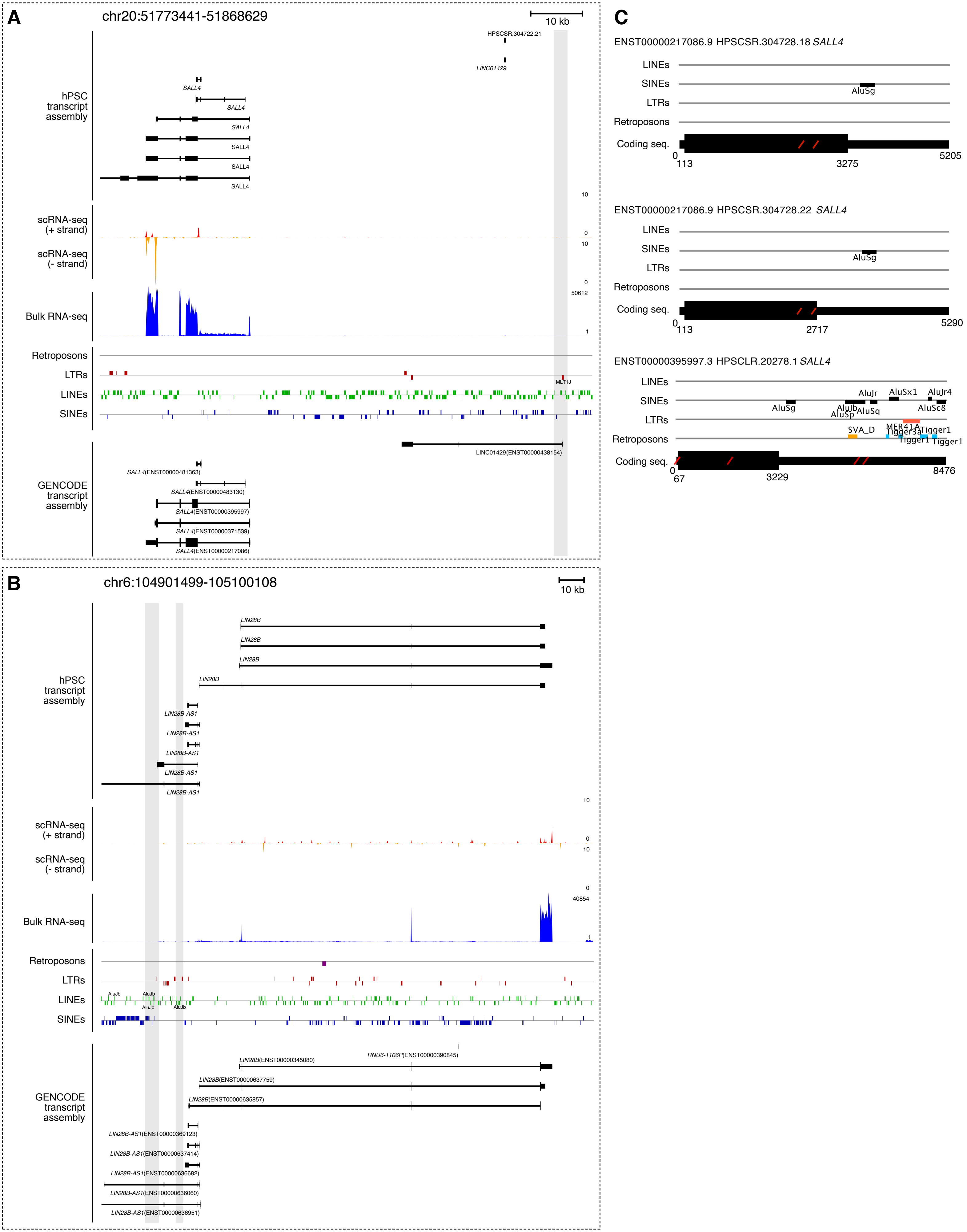
A
chr20:51773441-51868629
10 kb
hPSC transcript assembly
HPSCSR.304722.21
LINC01429
SALL4
scRNA-seq (+ strand)
scRNA-seq (- strand)
Bulk RNA-seq
Retroposons
LTRs
LINEs
SINEs
MLT1J
GENCODE transcript assembly
LINC01429(ENST00000438154)
SALL4(ENST00000481363)
SALL4(ENST00000483130)
SALL4(ENST00000395997)
SALL4(ENST00000371539)
SALL4(ENST00000217086)
B
chr6:104901499-105100108
10 kb
LIN28B
LIN28B-AS1
AluJb
RNU6-1106P(ENST00000390845)
LIN28B(ENST00000345080)
LIN28B(ENST00000637759)
LIN28B(ENST00000635857)
LIN28B-AS1(ENST00000369123)
LIN28B-AS1(ENST00000637414)
LIN28B-AS1(ENST00000636682)
LIN28B-AS1(ENST00000636060)
LIN28B-AS1(ENST00000636951)
C
ENST00000217086.9 HPSCSR.304728.18 SALL4
LINEs
SINEs
LTRs
Retroposons
Coding seq.
AluSg
0
113
3275
5205
ENST00000217086.9 HPSCSR.304728.22 SALL4
AluSg
0
113
2717
5290
ENST00000395997.3 HPSCLR.20278.1 SALL4
AluSg
AluJr
AluJb
AluSp
AluSq
AluSx1
AluJr4
AluSc8
SVA_D
MER41A
Tigger3a
Tigger1
0
67
3229
8476

### Figure S9
